# Automated compound testing in zebrafish xenografts identifies combined MCL-1 and BCL-X_L_ inhibition to be effective against Ewing sarcoma

**DOI:** 10.1101/2021.06.17.448794

**Authors:** Sarah Grissenberger, Caterina Sturtzel, Andrea Wenninger-Weinzierl, Eva Scheuringer, Lisa Bierbaumer, Susana Pascoal, Marcus Tötzl, Eleni Tomazou, Olivier Delattre, Didier Surdez, Heinrich Kovar, Florian Halbritter, Martin Distel

**Affiliations:** St. Anna Children’s Cancer Research Institute, Zimmermannplatz 10, 1090 Vienna, Austria; Zebrafish platform Austria for preclinical drug screening, Zimmermannplatz 10, 1090 Vienna, Austria; INSERM U830, Équipe Labellisée LNCC, Diversity and Plasticity of Childhood Tumors Lab, PSL Research University, SIREDO Oncology Centre, Institut Curie Research Centre, 75005 Paris, France; Balgrist University Hospital, University of Zurich, Zurich, Switzerland; Dept. Pediatrics, Medical University Vienna, Währinger Gürtel 18-20, 1090 Vienna, Austria

**Keywords:** Ewing sarcoma, zebrafish, xenotransplantation, compound screening, anti-apoptotic protein inhibition, high-content imaging

## Abstract

Ewing sarcoma is a pediatric bone and soft tissue cancer for which new therapies to improve disease outcome and to reduce adverse effects of current standard treatments are urgently needed. To identify new and effective drugs, phenotypic drug screening has proven to be a powerful method and a cancer model ideally suited for this approach is the larval zebrafish xenograft system. Complementing mouse xenografts, zebrafish offer high-througput screening possibilities in an intact complex vertebrate organism. Here, we generated Ewing sarcoma xenografts in zebrafish larvae and established a workflow for automated imaging of xenografts, tumor cell recognition within transplanted zebrafish and quantitative tumor size analysis over consecutive days by high-content imaging. The increased throughput of our *in vivo* screening setup allowed us to identify combination therapies effective against Ewing sarcoma cells. Especially, combined inhibition of MCL-1 and BCL-X_L_, two anti-apoptotic proteins, was highly efficient at eradicating tumor cells in our zebrafish xenograft assays with two Ewing sarcoma cell lines and with patient-derived cells. Transcriptional analysis across Ewing sarcoma cell lines and tumors revealed that *MCL-1* and *BCL2L1*, coding for BCL-X_L_, are the most abundantly expressed anti-apoptotic genes, suggesting that combined MCL-1/BCL-X_L_ inhibition might be a broadly applicable strategy for Ewing sarcoma treatment.

## Introduction

Ewing sarcoma is the second most frequent malignant bone and soft tissue tumor in children and adolescents, affecting around 1.5 per million children in populations of European descent. 20-25% of all cases show metastases at the time of diagnosis and another 30% relapse with distant metastases, which are often resistant to therapy, resulting in a low 5-year overall survival of <30% for those patients^1,2^.

Genetically, Ewing sarcoma is defined by a chromosomal translocation (most often t(11;22)(q24;q12)) leading to the expression of the aberrant transcription factor EWS-FLI1^3^. Despite this specific genetic aberration having been known for more than two decades, attempts to engineer a genetic animal model have failed so far^4^. The main obstacle is the still elusive cell of origin of Ewing sarcoma, which prevents targeted expression of EWS-FLI1 in the correct and permissive cell type *in vivo*^4^. The lack of a genetic animal model adequately mimicking the disease is hampering the development of new therapeutic strategies. Alternatively, xenotransplantation of cell line or patient-derived human Ewing sarcoma cells into immune compromised mice has been successfully applied to characterize Ewing sarcoma tumor cells *in vivo* and to identify novel treatment strategies^5-8^. However, mouse xenografts are laborious and costly to establish preventing their broad application in drug screening. In recent years, zebrafish (*Danio rerio*) tumor cell xenotransplantation has emerged as a complementary system to mouse xenografts^9-14^. In particular, xenografts in transparent zebrafish embryos and larvae offer an opportunity for *in vivo* live imaging of tumor cell proliferation and migration in an intact transparent organism. Furthermore, phenotypic drug testing is fast, technically easy, and cost-efficient in zebrafish xenograft models as small compounds can be administered directly into the water surrounding the larvae in 96-well format^15,16^. Ewing sarcoma cell lines have previously been transplanted into the yolk of zebrafish embryos and these xenografts have been applied to evaluate anti-tumor effects of selected compounds^17,18^.

In this study, we investigated the suitability of larval zebrafish Ewing sarcoma xenografts for automated high-througput *in vivo* compound testing. We show that Ewing sarcoma cells transplanted into the perivitelline space of zebrafish larvae, a xenograft site alternative to the yolk, persist, proliferate and form vascularized aggregates resembling small tumors. Furthermore, we established a workflow for automated image acquisition, tumor recognition and tumor size quantification of xenografted zebrafish in 96-well format. Our setup allowed us to test the *in vivo* efficacy of compounds with previously reported *in vitro* activity against Ewing sarcoma cells. We were able to identify combination treatments, which showed enhanced efficacy in our xenograft model. Especially, combining two anti-apoptotic protein inhibitors, S63845 (inhibiting MCL-1) and A-1331852 (inhibiting BCL-X_L_), could efficiently eliminate tumor cells. Efficacy of this drug combination was also confirmed on zebrafish xenografts with a second cell line and patient-derived Ewing sarcoma cells. Furthermore, transcriptional analysis revealed that MCL-1 and BCL-X_L_ are anti-apoptotic proteins broadly expressed across Ewing sarcoma cell lines and tumors, suggesting that their combined inhibition promises to be a widely applicable strategy for Ewing sarcoma treatment.

## Results

### Ewing sarcoma cells survive and proliferate in zebrafish xenografts

In order to establish Ewing sarcoma xenografts in zebrafish, we first investigated if SK-N-MC cells can survive and proliferate in zebrafish embryos upon xenotransplantation at 2 days post fertilization (dpf). We used a GFP expressing sub-clone (shSK-E17T) that carries a doxycycline-inducible shRNA to EWS-FLI1, allowing for modulation of the driver oncogene^19^. We initially compared two common injection sites, yolk and perivitelline space (PVS) and observed consistent growth of shSK-E17T cells in the PVS, whereas cell numbers decreased in the yolk already after 24 hours post injection (hpi) (Fig. S1). Thus, we applied PVS injection for all subsequent xenotransplantation experiments. At 2 hpi, we found tumor cells to be dispersed at the injection site, but to form a compact mass by 1 day post injection (dpi), with an increase in size over subsequent days (Fig. 1A). Ki67 and activated Caspase 3 immunofluorescence staining from 2 hpi up to 7 dpi confirmed that shSK-E17T proliferate in zebrafish embryos/larvae and that only few tumor cells undergo apoptosis (Fig. 1B-C, Fig. S2). This indicated that shSK-E17T are a suitable tumor cell line for establishing Ewing sarcoma xenografts in zebrafish.

**Figure 1:**
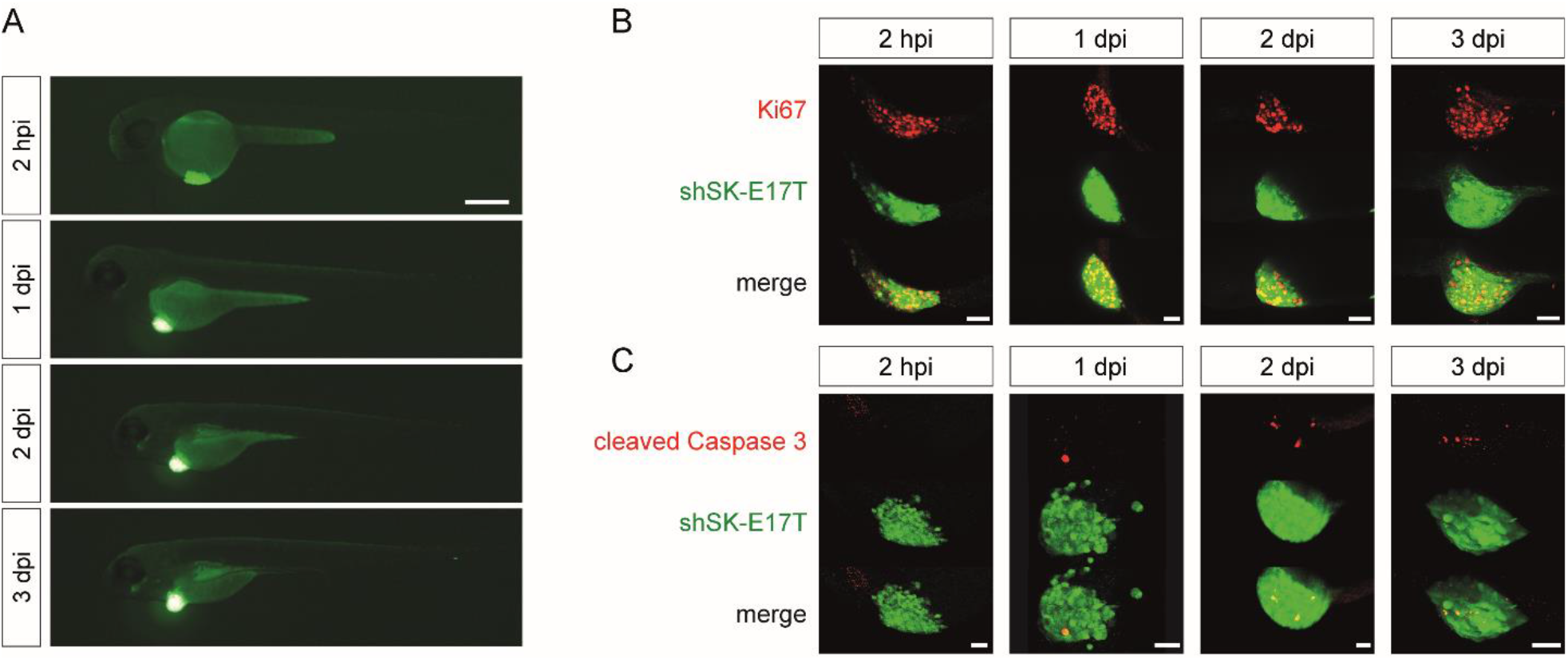
shSK-E17T Ewing sarcoma zebrafish xenografts. A) Zebrafish xenografted with GFP-expressing shSK-E17T Ewing sarcoma cells (green) imaged on a fluorescence microscope over consecutive days from 2 hpi to 3 dpi. B) Immunostaining for human Ki67 revealing proliferating tumor cells and C) for cleaved Caspase 3 demarcating apoptotic cells at different time points from 2 hpi to 3 dpi. (B, n: 2 hpi = 5, 1 dpi = 7, 2 dpi = 5, 3 dpi = 6); (C, n: 2 hpi = 6, 1 dpi = 8, 2 dpi = 5, 3 dpi = 7). Scale bars are 250 μm in A and 50 μm in B and C.

### Establishing automated imaging and quantification of zebrafish xenografts

In order to screen small compounds for tumor-growth-inhibiting effects at medium to high-througput, we next sought to automate image acquisition and image analysis of xenografted zebrafish larvae enabling reproducible quantification of tumor sizes at different time points. We applied a high-content imager (Operetta CLS, PerkinElmer) for automated image acquisition of xenografted zebrafish larvae in 96-well format. To ensure proper positioning for lateral imaging and to minimize imaging time, we used 96-well imaging plates with predefined slots for zebrafish larvae (ZF plates, Hashimoto) (Fig. 2A).

**Figure 2:**
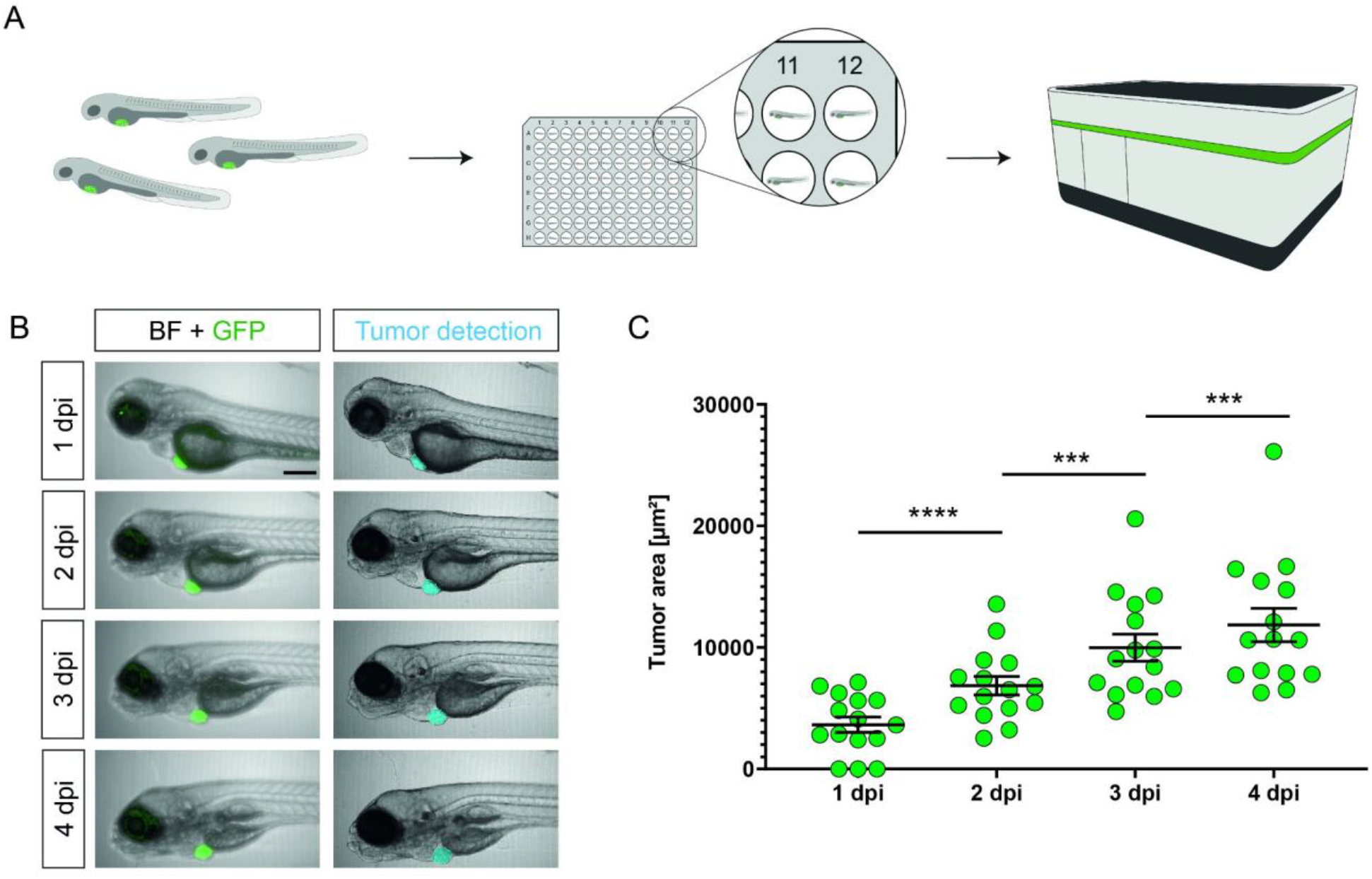
Automated imaging and tumor size quantification of xenotransplanted zebrafish larvae. A) Schematic representation of the imaging workflow: Xenotransplanted larvae were anesthetized, embedded in 96-well ZF plates (1 larva/well) and imaged in brightfield (BF) and fluorescence on the Operetta high-content imager (PerkinElmer). B) Representative images of shSK-E17T zebrafish xenografts imaged on consecutive days (1 dpi - 4 dpi), automated tumor detection and tumor area quantification. C) Measured tumor size areas from 1 dpi to 4 dpi (n = 15). Scale bar in B is 250 μm. Statistical analysis in C was performed with a repeated measure (RM) one-way ANOVA, ****: p≤0.0001, ***: p≤0.001. Error bars represent SEM of 15 individual larvae.

This setup allowed us to record images of anesthetized xenografted larvae in bright field and fluorescence at higher throughput once per day for 4 consecutive days from 1 dpi to 4 dpi (n = 15) (Fig. 2B). Typical imaging time for one plate with 96 larvae was around 60 minutes (23 planes per larva, bright field and one fluorescent channel in confocal mode).

To automate tumor cell detection and quantification of the size of the tumor cell cluster, we customized an analysis module within the Harmony software (PerkinElmer) for our specific application in zebrafish. We found that footprint projection to measure the tumor area delivered the most precise and reproducible results across many xenografted zebrafish. Consistent with our Ki67 immunostaining results, measuring the tumor size with this analysis module revealed a significant increase from 1 dpi to 4 dpi (RM one-way ANOVA, p ≤ 0.0001), but also between each measured time point (p ≤ 0.001 throughout), confirming that tumor cells are proliferating in our zebrafish xenografts (Fig. 2B-C).

### Automated small compound testing in Ewing sarcoma zebrafish xenografts

Having established a xenograft model, an imaging setup and an analysis method, which enabled us to quantify tumor sizes over several consecutive days in an automated way prompted us to apply this assay for screening small compounds for anti-tumor activity in Ewing sarcoma. Here, we decided to focus on compounds, which had previously shown *in vitro* efficacy against Ewing sarcoma cell lines. We also included compounds, which are clinically applied like doxorubicin or irinotecan and we added venetoclax to investigate the role of different anti-apoptotic proteins. In total, we tested 16 compounds (see Table S1).

We first identified a maximum concentration for each compound without any apparent adverse morphological effect, like edema, malformations, or lethality (Table S1). This “no observed effect concentration” (NOEC) was determined by treating zebrafish larvae incubated from approx. 76 hpf to 5 dpf to best mimic the actual drug treatment period of xenografted zebrafish. Next, we tested all 16 compounds at maximum NOEC on xenografted zebrafish for their *in vivo* anti-Ewing-sarcoma efficacy. For this, shSK-E17T cells were injected into zebrafish embryos at 48 hpf. An image of the xenografted larvae was recorded at 1 dpi on the high-content imager. Subsequently, the respective compounds were administered to the water surrounding the xenografted larvae. Larvae were incubated in compounds or in respective concentrations of DMSO for 2 days and an image of each larva was again recorded at 3 dpi (Fig. 3A). The tumor area was quantified and its size at 3 dpi was compared to the size at 1 dpi for each treated larva.

**Figure 3:**
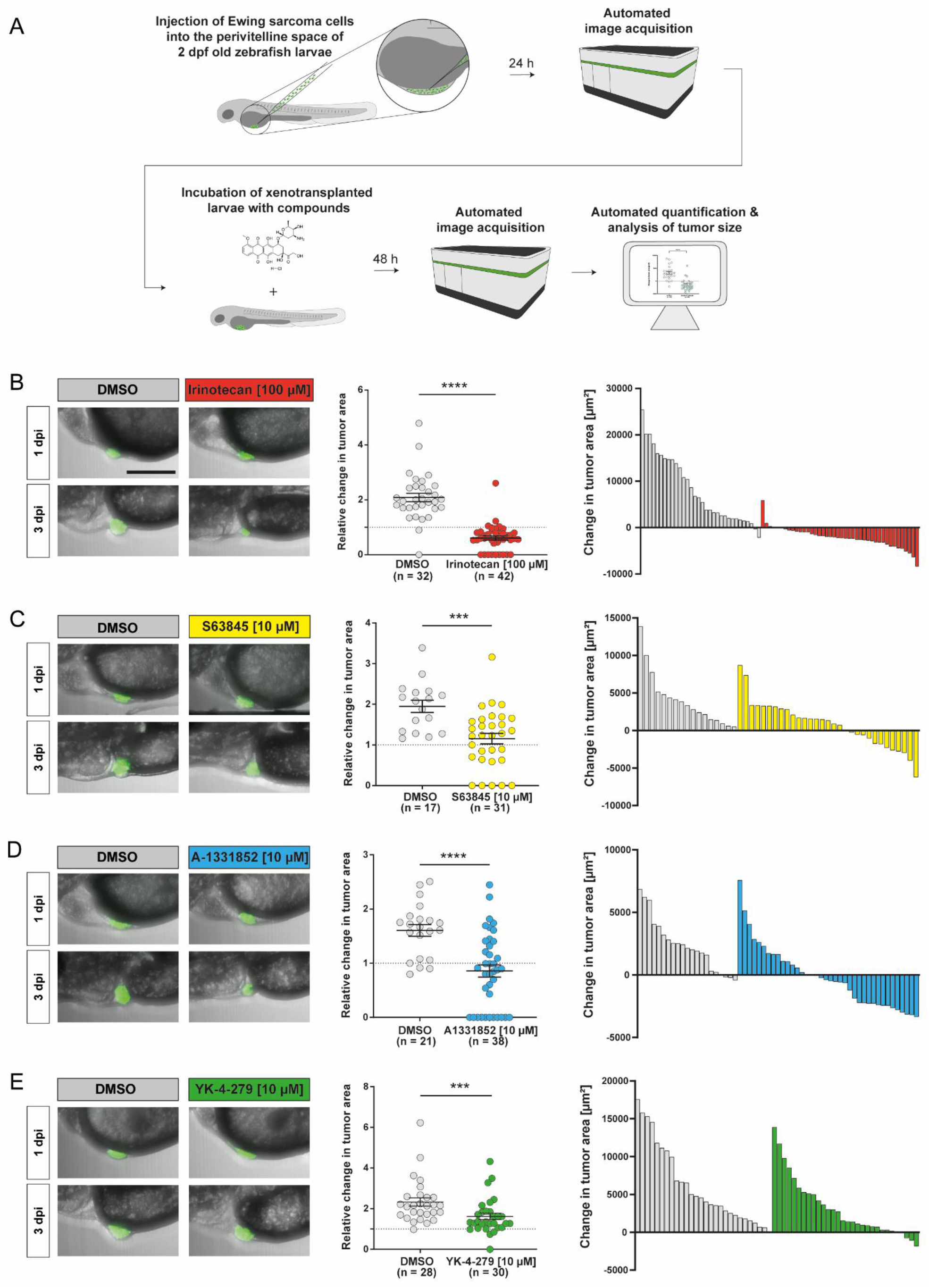
Single agent treatment of xenotransplanted larvae. A) Schematic representation of the workflow - Zebrafish embryos were xenotransplanted with shSK-E17T Ewing sarcoma cells at 2 dpf. After 24 hours (1 dpi) embryos were imaged on the Operetta CLS and subsequently incubated in respective amounts of small compounds. After 48 hours (3 dpi) larvae were imaged again and changes in tumor size were analyzed. B-E) Xenotransplanted larvae were treated with irinotecan (100 μM, B), S63845 (10 μM, C), A-1331852 (10 μM, D) and YK-4-279 (10 μM, E) for 48 hours (1 dpi - 3 dpi). Dot plots show relative changes of tumor area (3 dpi/1 dpi). Waterfall plots (right panels, one bar per zebrafish) total change in tumor area (3 dpi - 1 dpi). Scale bar is 250 μm. Statistical analyses were performed with a Mann-Whitney test, ****: p≤0.0001, ***: p≤0.001. Error bars represent SEM of combined larvae from two independent experiments.

Out of the 16 tested compounds, 4 showed an effect on tumor growth. A strong decrease in tumor size was observed upon treatment with the topoisomerase I inhibitor irinotecan (−35% on 3 dpi compared to 1 dpi; control group 123% increase) (Fig. 3B). Furthermore, treatment with the MCL-1 inhibitor S63845 (10 μM) reduced tumor growth to 18% (control group 96% increase) (Fig. 3C). The BCL-X_L_ inhibitor A-1331852 (10 μM) reduced tumor size by −5% (control group 65% increase) (Fig. 3D). YK-4-279, an inhibitor reported to block the interaction between EWS-FLI1 and RNA helicase A decreased tumor growth to 62% (control group 134% increase) (Fig. 3E)^20,21^.

Titration experiments with S63845, A-1331852, YK-4-279, and irinotecan confirmed that the observed effects on tumor growth were dose-dependent. All other tested compounds did not show any significant effect in our *in vivo* model (Fig. S3).

### Identification of effective combination treatments

We next investigated combinations of compounds with single agent efficacy (S63845, A-1331852, YK-4-279 and irinotecan) for potential synergistic effects. First, we determined NOECs for all possible combinations (Table S2) and then tested compound combinations at maximum NOEC on shSK-E17T zebrafish xenografts following the same workflow as for single compound treatments. While combining YK-4-279 (5 μM) with irinotecan (5 μM) or A-1331852 (10 μM) had no additional effect (Fig. S4), a combination of YK-4-279 (5 μM) and S63845 (5 μM) could further reduce tumor size (−31.1 %) than single compounds alone (DMSO control: 107.7%; S63845 (5 μM) 44.1 %; YK-4-279 (5 μM) 35 %) (Fig. 4). The combinations of irinotecan (5 μM) with either S63845 (5 μM) or A-1331852 (10 μM) showed strong enhanced average reduction of tumor size (irinotecan (5 μM)/S63845 (5 μM) −45.8%; irinotecan (5 μM)/A-1331852 (10 μM) −49 %) (Fig. 4). Combining S63845 (5 μM) with A-1331852 (10 μM) was able to fully eradicate Ewing sarcoma tumor cells in our xenograft setting with no apparent adverse effects (Fig. 4). Dual inhibition of MCL-1 and BCL-X_L_ was thus identified as a powerful strategy for Ewing sarcoma treatment in our model.

**Figure 4:**
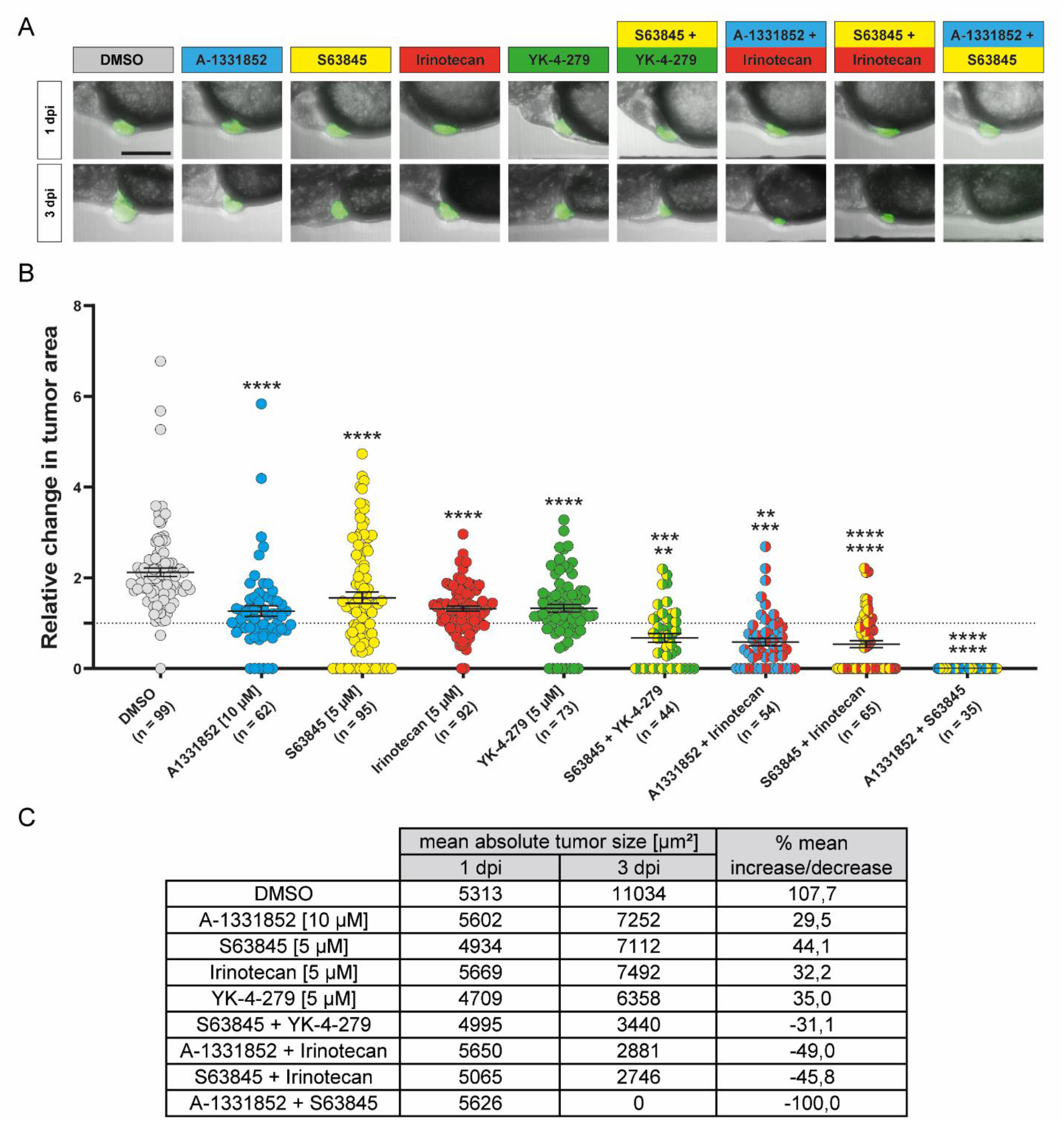
Combination treatments of xenotransplanted larvae. Xenotransplanted larvae were treated with A-1331852 (10 μM), S63845 (5 μM), Irinotecan (5 μM), YK-4-279 (5 μM) and respective combinations for 48 hours (1 dpi - 3 dpi). A) Representative images for average change in tumor size for every condition. B) Relative change of tumor area (3 dpi/1 dpi). C) Mean absolute tumor sizes and percent changes. Scale bar is 250 μm. Significance is shown as followed: Single treatments were compared to DMSO control. Combination treatments were compared to respective single treatments (top value indicates comparison to first compound, bottom value indicates comparison to second compound). Statistical analyses were performed with a Kruskal-Wallis test, ****: p≤0.0001, ***: p≤0.001, **: p≤0.01. Error bars represent SEM of combined larvae from two independent experiments.

### Combination treatments are able to eradicate Ewing sarcoma cells with lowered EWS-FLI1 expression levels

Heterogeneity among tumor cells is one factor underlying selective responses to therapy. Fluctuations in EWS-FLI1 levels have been postulated to contribute to Ewing sarcoma plasticity in patients and it has been demonstrated that *in vitro* modulation of EWS-FLI1 levels can switch Ewing sarcoma cells from a proliferative to a migratory/invasive phenotype with enhanced metastatic potential *in vivo^19,22^*. Cells expressing low levels of EWS-FLI1 are believed to contribute to resistance to standard therapy and might be responsible for relapse as they can switch back to the more proliferative state leading to tumor colonization at metastatic sites. In order to investigate, if any of the four effective compound combinations would also target EWS-FLI1^low^ cells, we took advantage of the inducible EWS-FLI1 knockdown system in shSK-E17T cells^19^. Upon treatment with doxycycline, shSK-E17T cells expressed a small hairpin (sh)RNA targeted against the EWS-FLI1 transcript leading to ~42% knockdown of the fusion protein (Fig. S5A). Indeed, we observed a significant decrease in tumor growth from 1 dpi to 3 dpi upon EWS-FLI1 knockdown compared to control xenografts (Fig. S5B), which is in line with a decrease in proliferation under lower EWS-FLI1 levels. An increase in migration was however not observed during our 4 day assay. In this xenograft setting with reduced EWS-FLI1 levels, the combination of YK-4-279 (5 μM) and S63845 (5 μM) could not reduce tumor size further than YK-4-279 (5 μM) alone. However, irinotecan (5 μM) with S63845 (5 μM) or with A-1331852 (10 μM) still showed increased efficacy over single compound treatment. Finally, the combination of S63845 (5 μM) with A-1331852 (10 μM) remained most effective, eradicating all tumor cells in 26 out of 32 xenografts (Fig. 5). Thus, also under reduced EWS-FLI1 levels, dual MCL-1 and BCL-X_L_ inhibition is highly effective in our model.

**Figure 5:**
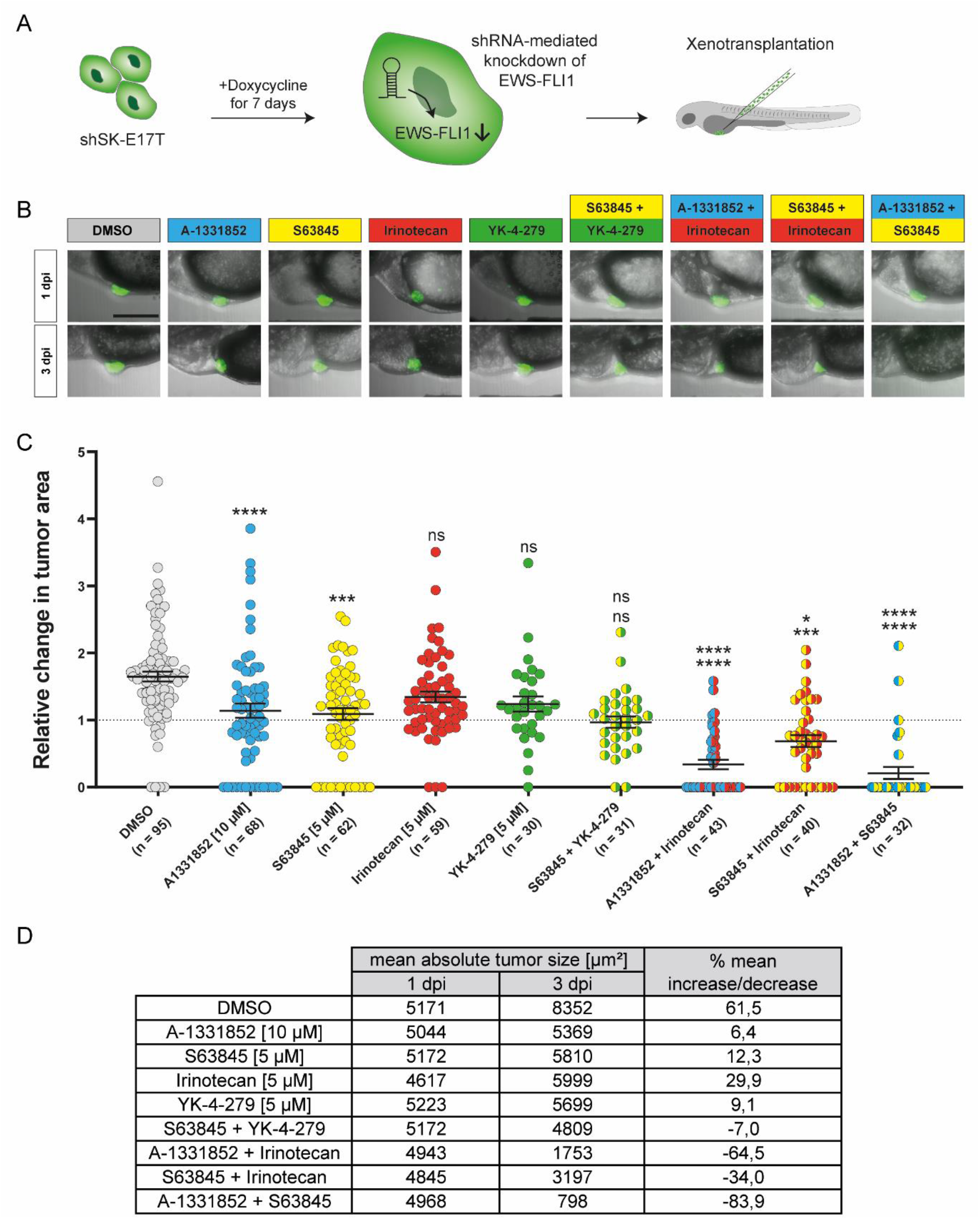
Combination treatments of xenotransplanted larvae in EWS-FLI1^low^ conditions. A) Schematic representation of the workflow: shSK-E17T cells were pre-treated with doxycycline for 7 days in culture to induce knockdown of EWS-FLI1. Xenotransplanted embryos were incubated in E3 with doxycycline over the course of the experiment. Embryos were treated with A-1331852 (10 μM), S63845 (5 μM), irinotecan (5 μM), YK-4-279 (5 μM) and respective combinations for 48 hours (1 dpi - 3 dpi). B) Representative images for average change in tumor size for every condition. C) Relative change of tumor area (3 dpi/1 dpi). D) Mean absolute tumor sizes and percent changes. Scale bar is 250 μm. Significance is shown as followed: Single treatments were compared to DMSO control. Combination treatments were compared to respective single treatments (top value indicates comparison to first compound, bottom value indicates comparison to second compound). Statistical analyses were performed with a Kruskal-Wallis test, ****: p≤0.0001, ***: p≤0.001, *: p≤0.05. Error bars represent SEM of combined larvae of two independent experiments.

### Validation of combination treatments

In order to investigate whether the vulnerability towards our identified combination treatments was specific to the shSK-E17T cell line, we repeated combination treatments on zebrafish xenografted with a second Ewing sarcoma cell line, A673-1c, modified to express tagRFP for visualization on the high-content imager^19^. Similar to shSK-E17T xenografts, A673-1c cells formed tumors, which increased in size over 3 days. However, A673-1c proliferated less with a mean 1.45-fold increase in tumor size from 1 dpi to 3 dpi compared to a 2.1-fold increase of shSK-E17T cells. All three combination treatments were able to reduce tumor size in A673-1c xenografts and again MCL-1 and BCL-X_L_ dual inhibition eradicated all tumor cells confirming our previous results (Fig. S6).

We next also aimed at testing the most promising combination on Ewing sarcoma xenografts with patient-derived cells in zebrafish. For this, we established xenografts from patient-derived cells, which had been propagated in mouse PDXs^7,8^. After dissociation of tumor pieces and short-term *in vitro* culture, patient-derived cells were labeled with CellTrace™ Violet (ThermoFisher Scientific) and transplanted into 2 dpf old zebrafish embryos. Patient-derived tumor cells persisted in xenografted zebrafish over 3 days and we observed a 1.45-fold increase in tumor size from 1 dpi to 3 dpi comparable to A673-1c cells (Fig. 6). Co-treatment with S63845 (5 μM) and A-1331852 (10 μM) fully eliminated all tumor cells in 55 out of 57 zPDXs confirming our previous results and suggesting a potential broad efficacy of this drug combination against Ewing sarcoma (Fig. 6).

**Figure 6:**
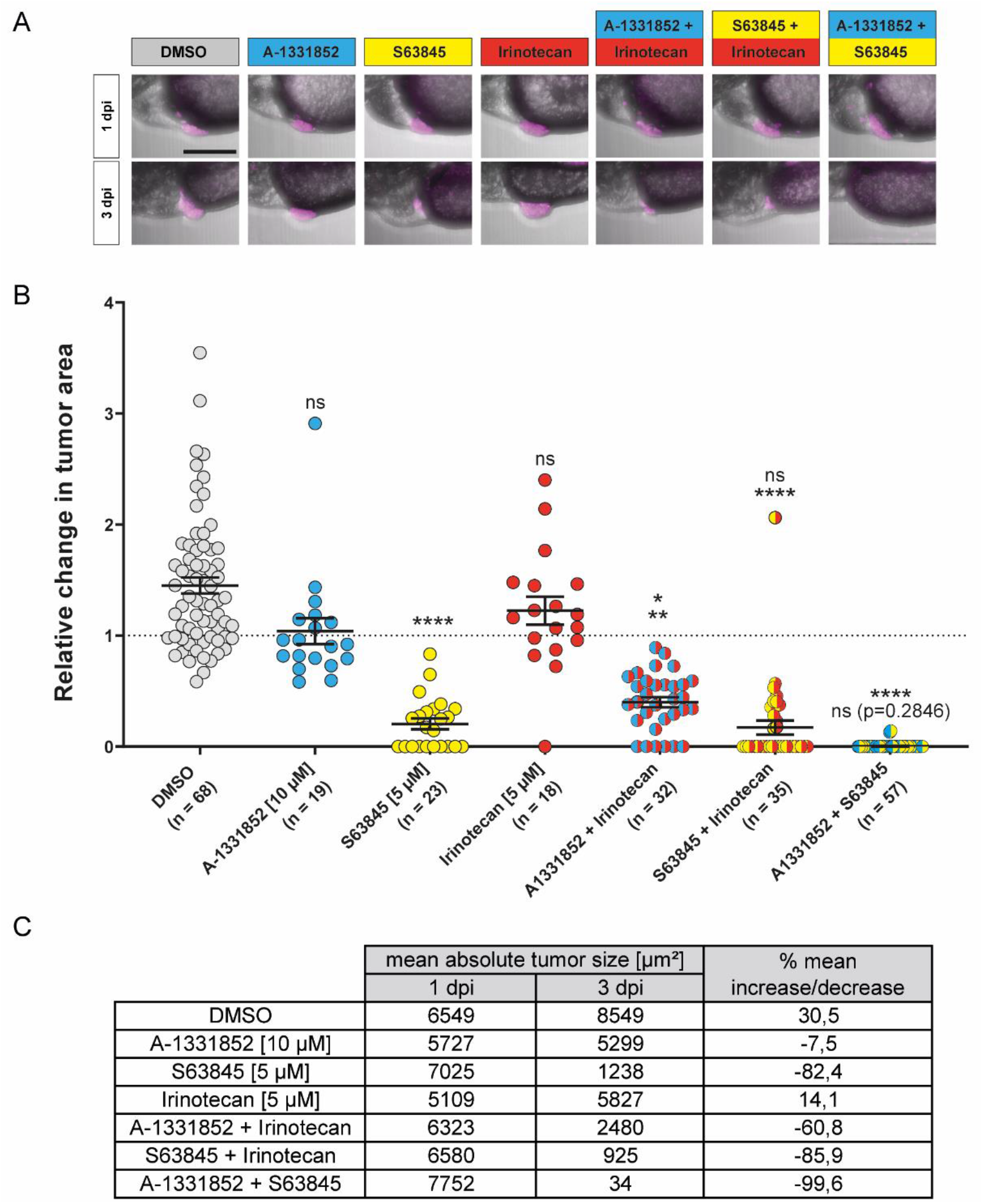
Combination treatments in patient-derived cells. Xenotransplanted larvae were treated with A-1331852 (10 μM), S63845 (5 μM), irinotecan (5 μM), and respective combinations for 48 hours (1 dpi - 3 dpi). A) Representative images for average change in tumor size for every condition. B) Relative change in tumor area (3 dpi/1 dpi). C) Mean absolute tumor sizes and percent changes. Scale bar is 250 μm. Significance is shown as followed: Single treatments were compared to DMSO control. Combination treatments were compared to respective single treatments (top value indicates comparison to first compound, bottom value indicates comparison to second compound). Statistical analyses were performed with a Kruskal-Wallis test, ****: p≤0.0001, **: p≤0.01, *: p≤0.05. Error bars represent SEM of combined larvae (n) of one experiment (single treatments) or two experiments (combinations).

### MCL-1/BCL-X_L_ dual inhibition as treatment strategy in Ewing sarcoma

In order to gauge applicability of dual MCL-1 and BCL-X_L_ inhibition for Ewing sarcoma treatment in general, we interrogated gene expression data from Ewing sarcoma tumors and cell lines obtained from the Treehouse Childhood Cancer Initiative^23^. In addition to *MCL1* and *BCL2L1* (encoding BCL-X_L_), we checked the expression levels of the related anti-apoptotic genes *BCL2, BCL2L2* (encoding BCL-w), and *BCL2A1*, as well as the major known effector genes *BAK, BAX* and *BOK*. We found consistent expression of *MCL1, BCL2L1* and the major effector gene *BAX* across Ewing sarcoma samples and cell lines (Fig. 7). In contrast, other anti-apoptotic genes (*BCL2, BCL2A1*) and effector genes (*BOK*) were only detected at lower mRNA levels. In line with our experimental data, this suggested MCL-1 and BCL-X_L_ to be major anti-apoptotic players in Ewing sarcoma, providing a rationale for their combined inhibition as therapeutic strategy.

**Figure 7:**
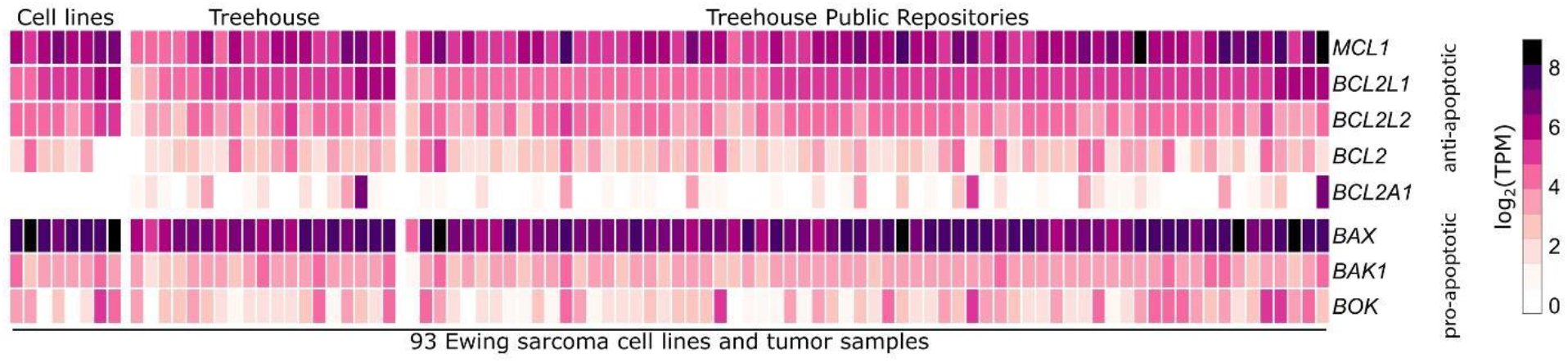
Expression analysis of anti- and pro-apoptotic genes across Ewing sarcoma samples. Heatmap displaying normalized gene expression levels of anti-apoptotic genes *MCL1, BCL2L1, BCL2L2, BCL2*, and *BCL2A1* as well as pro-apoptotic effectors *BAX, BAK1*, and *BOK*. Expression was analyzed in RNA-seq data across Ewing sarcoma cell lines (A-673, TC-71 CADO-ES1, SK-ES-1, RD-ES, MHH-ES-1 Hs 863.T and HS 822.T) and tumors. RNA-seq data were obtained from the Treehouse Childhood Cancer Initiative^23^. TPM, transcripts per million.

## Discussion

Larval zebrafish xenografts are gaining increasing attention in cancer research as a vertebrate model system complementary to mouse xenografts^24^. Due to their small size, transparency and ease of compound administration, zebrafish larvae are well suited for small compound screening^16^. Although, a proof-of-concept study demonstrated that high-content imaging can be applied for phenotype-based compound testing on zebrafish xenografts with leukemia cells (K562) injected into the yolk^25^, this approach has not been widely adopted subsequently and the potential of zebrafish xenografts for the discovery of new therapeutic compounds and in particular of effective compound combinations by automated high-throughput high-content screening has not been fully exploited yet.

In this study, we generated Ewing sarcoma xenografts from two cell lines and patient-derived cells in zebrafish larvae and applied them as *in vivo* models for small compound screening. To increase the drug screening throughput, we established an automated workflow for image acquisition of xenografted zebrafish, tumor cell detection, and tumor size quantification using a high-content imager (Operetta CLS, PerkinElmer). The increased throughput achieved through this automation merited systematic analysis of more than 3,500 xenotransplanted zebrafish larvae in total and identified 3 drug combinations with high efficacy against Ewing sarcoma cells. With our current setup, around 12 small compounds can be tested on approximately 20 xenografted zebrafish larvae per week in an academic setting – a throughput not achievable by most labs in mammalian xenograft models.

Out of the 15 compounds with previously reported *in vitro* efficacy against Ewing sarcoma cells tested in our assay, we confirmed significant *in vivo* effects for irinotecan, S63845, A-1331852 and YK-4-279. The topoisomerase I inhibitor irinotecan has been investigated in several clinical trials e.g. together with temozolomide for Ewing sarcoma (see https://clinicaltrials.gov/ct2/results?cond=Ewing+Sarcoma&term=irinotecan&cntry=&state=&city=&dist=)^26^. A variant of YK-4-279 with improved bioavailability (TK216) has also entered clinical phase I trials (https://clinicaltrials.gov/ct2/show/NCT02657005?term=tk216&rank=1). However, to date, efficacy against Ewing sarcoma cells of two anti-apoptotic protein inhibitors, S63845 directed against MCL-1 and A-1331852 targeting BCL-X_L_ had only been reported *in vitro^27^*.

Inhibition of anti-apoptotic proteins is emerging as a promising strategy for cancer treatment. The tremendous success of the BCL-2 specific inhibitor venetoclax in the treatment of chronic lymphocytic leukemia encouraged the development of inhibitors for other anti-apoptotic proteins of the BCL-2 family like MCL-1 or BCL-X_L_^28^. Cancer cells seem to be more primed for apoptosis than most healthy cells and often depend on anti-apoptotic proteins for survival^29^. In Ewing sarcoma, MCL-1 has been identified as a dependency protein in CRISPR screens, which is in good agreement with our drug testing results^30,31^.

We show that combining irinotecan with either the MCL-1 inhibitor S63845 or the recently developed BCL-X_L_ inhibitor A-1331852 shows increased efficacy over single agent treatment^32^. To our knowledge, S63845 has not been applied together with irinotecan for cancer treatment, but enhanced efficacy of irinotecan when combined with A-1331852 was reported in colorectal cancer xenografts in mouse^32^. A possible explanation for this effect is that most chemotherapeutics like DNA-damaging agents or other cytotoxic drugs produce cellular stress that leads to upregulation of pro-apoptotic proteins priming the cell for apoptosis, but are kept in check by anti-apoptotic proteins^29^. Inhibition of the relevant anti-apoptotic proteins may release this break.

The most effective combination eradicating almost all tumor cells in our zebrafish xenografts with two Ewing sarcoma cell lines as well as with patient-derived Ewing sarcoma cells was dual MCL-1 and BCL-X_L_ inhibition with S63845 and A-1331852. Interestingly, high expression of BCL-X_L_ has been reported as a predictive factor for MCL-1 inhibition resistance, indicating that BCL-X_L_ can functionally compensate for MCL-1^33^. Our gene expression analysis revealed that next to *MCL-1, BCL2L1* (BCL-X_L_) is highly expressed across Ewing sarcoma samples. In good agreement with our *in vivo* results, exquisite sensitivity to combined MCL-1/BCL-X_L_ inhibition was recently also reported for a larger panel of solid pediatric tumor cell lines *in vitro*, including two Ewing sarcoma cell lines, SK-ES-1 and A4573^27^.

We also noticed different sensitivities for MCL-1 and BCL-X_L_ inhibition in our three models. Xenografts with patient-derived cells were highly sensitive to MCL-1 inhibition (Fig. 6), whereas A673 xenografts showed sensitivity to BCL-X_L_ inhibition (Fig. S6). This suggests, that dual MCL-1/BCL-X_L_ inhibition will cover this range of differing sensitivities to single compounds of Ewing sarcoma cells. Furthermore, shSK-E17T cells remained responsive to combined MCL-1/BCL-X_L_ inhibition in xenografts upon knockdown of EWS-FLI1, indicating that lower EWS-FLI1 levels, as might occur during metastasis, do not lead to treatment escape of Ewing sarcoma cells. Together, this provides a rationale for dual inhibition of MCL-1 and BCL-X_L_ for Ewing sarcoma treatment. So far, promising preclinical *in vivo* data for MCL-1/BCL-X_L_ inhibition strategy is available for difficult-to-treat melanoma in mouse xenografts with tolerable toxicity and for rhabdomyosarcoma using Kym-1 cells transplanted onto the chorioallantoic membrane of chicken eggs^27,34^.

Although, anti-apoptotic protein inhibitors are in clinical trials for many adult tumor entities, their application for pediatric cancers needs very careful evaluation of potentially occurring toxicities due to cell death of normal tissues. It was previously reported that hematopoietic organs in adults are primed for apoptosis and hence show sensitivity to anti-apoptotic protein inhibition^35^. For example, thrombocytopenia is a known effect of systemic BCL-X_L_ inhibition^36^. In contrast, cells in the brain are refractory to apoptosis in adults, but they seem to be sensitive at early developmental stages^35^. Nevertheless, BH3 profiling revealed that cells in the human brain transition from being primed for apoptosis to full apoptotic resistance around approximately 6 years of age^35^. This indicates that potential brain toxicity might not preclude application of anti-apoptotic protein inhibitors in older Ewing sarcoma patients.

We did not observe adverse effects on zebrafish health in our short-term assay with anti-apoptotic protein inhibitors S63845 and A-1331852 as single agents (10 μM) or in combination (S63845 5 μM; A-1331852 10 μM). However, from mouse studies it is known that S63845 has six-fold higher affinity to human MCL-1 compared to murine MCL-1, hence observed toxicities vary between these species. Currently, it is unclear, if and how much S63845 affinities differ for human MCL-1 and its two zebrafish orthologs Mcl-1a and Mcl-1b and A-1331852 affinity for zebrafish BCL-X_L_ is also unknown. From our own analysis, we observed adverse effects of S63845 and A-1331852 at concentrations >10 μM in zebrafish larvae. Furthermore, S63845 showed efficacy in a zebrafish acute lymphoblastic leukemia model, clearly demonstrating that it does act on zebrafish MCL-1 orthologs^37^.

In conclusion, we present a setup for high-content imaging-assisted compound screening on zebrafish Ewing sarcoma xenografts, which identified combined MCL-1 and BCL-X_L_ inhibition to be highly effective against Ewing sarcoma cells. We anticipate that potential toxicities including thrombocytopenia can be overcome by dosing and timing as previously suggested^29,38^ and that dual MCL-1/BCL-X_L_ inhibition should be further investigated towards potential clinical application in Ewing sarcoma. The zebrafish xenograft model together with automated imaging and image analysis as presented here promises to be a powerful system widely applicable for drug testing for human tumor entities bridging the gap between *in vitro* screening and mouse xenografts. Furthermore, by using patient-derived zebrafish xenografts^9^ this setup can be adapted for personalized medicine approaches to identify patient-tailored drug combinations within a clinically relevant timespan.

## Materials & Methods

### Zebrafish strains and husbandry

Zebrafish (*Danio rerio*) were maintained at standard conditions^39,40^ according to the guidelines of the local authorities under licenses GZ:565304-2014-6 and GZ:534619-2014-4. For xenotransplantation experiments transparent zebrafish mutants (mitfa^b692/b692^; ednrba^b140/b140^) were used and experiments were performed under license GZ:333989-2020-4.

### Cell culture

GFP expressing shSK-E17T cells^19^ were cultured in RPMI 1640 medium with GlutaMAX™ (Gibco, Thermo Fisher Scientific, USA) supplemented with 10% FBS and 1% Penicillin-Streptomycin (10,000 U/ml, Gibco, thermos Fisher Scientific, USA) on fibronectin-coated plates. Knockdown of EWS-FLI1 in shSK-E17T cells was performed by shRNA induction by addition of 1 μg/ml Doxycycline (Sigma-Aldrich, USA) for 7 days pre-transplantation.

TagRFP expressing A673-1c cells^19^ were cultured in DMEM with GlutaMAX^TM^ (Gibco, Thermo Fisher Scientific, USA) supplemented with 10% FBS, 1% Penicillin-Streptomycin, 10 μg/ml Balsticidin (InvivoGen, USA) and 50 μg/ml Zeocin (InvivoGen, USA).

Primary Ewing sarcoma cells were obtained from a mouse patient-derived xenograft (IC-pPDX-87). Animal care and use for this study were performed in accordance with the recommendations of the European Community (2010/63/UE) for the care and use of laboratory animals. Experimental procedures were specifically approved by the ethics committee of the Institut Curie CEEA-IC #118 (Authorization APAFIS#11206-2017090816044613-v2 given by National Authority) in compliance with the international guidelines. Written informed consent allowing generation of PDX models was obtained.

Tumor pieces were dissociated according to Stewart et al. and PDX-87 cells were short-term cultured (<5 passages) in DMEM/F-12 with GlutaMAX™ (Gibco, Thermo Fisher Scientific, USA) supplemented with 1% B-27 (50X, Gibco, Thermo Fisher Scientific, USA) and 1% Penicillin-Streptomycin^41^. All cells were kept in an atmosphere with 5% CO_2_ at 37°C. At 90% confluence cells were passaged using Accutase (Pan Biotech, Germany).

### Preparation of cells for transplantation

To prepare shSK-E17T and A673-1c Ewing sarcoma cells for xenotransplantation cells were harvested with Accutase. After centrifugation (5 min, 500 g, 4°C) the pellet was resuspended in PBS supplemented with 2% FBS and put through a 35 μm cell strainer (5 ml polystyrene round bottom tubes with cell strainer cap, Corning, USA) to remove cell clumps. The total cell number was determined using a Coulter Counter (Z2, Beckman Coulter, USA). Cells were then centrifuged again, resuspended to a concentration of 100 cells/nl in PBS and kept on ice until transplantation.

To prepare PDX-87 cells for xenotransplantation cells were harvested with Accutase and counted. After centrifugation cells were resuspended at a concentration of 1*10^6^ cells/ml in PBS. CellTrace™ Violet (Invitrogen, Thermo Fisher Scientific, USA) was added to a final concentration of 5 μM. Cells were incubated with CellTrace™ Violet for 10 minutes at 37°C in the dark. 5 volumes of DMEM supplemented with 10% FBS were added and suspension was incubated for additional 5 min. After centrifugation (5 min, 500 g, 4°C) cells were resuspended in fresh DMEM supplemented with 10% FBS and incubated for 10 minutes in the dark. Finally, cells were put through a 35 μm cell strainer and total cell number was determined. Cells were then centrifuged again, resuspended to a concentration of 100 cells/nl in PBS and kept on ice until transplantation.

### Xenotransplantation

mitfa^b692/b692^; ednrba^b140/b140^ embryos were raised until 2 dpf at 28°C, dechorionated and anesthetized with 1x Tricaine (0.16 g/l Tricaine (Cat No. E1052110G, Sigma-Aldrich Chemie GmbH, Germany), adjusted to pH 7 with 1M Tris pH 9.5, in E3). For transplantation anesthetized larvae were aligned on a petri dish lid coated with a solidified 2% agarose solution as described previously^42^.

For injection of tumor cells, borosilicate glass capillaries (GB100T-8P, without filament, Science Products GmbH, Germany) pulled with a needle puller (P-97, Sutter Instruments, USA) were used. Needles were loaded with ~5 μl of tumor cell suspension, mounted onto a micromanipulator (M3301R, World Precision Instruments Inc., Germany) and connected to a microinjector (FemtoJet 4i, Eppendorf, Germany). Ewing sarcoma cells were injected into the perivitelline space (PVS) of zebrafish larvae. Xenotransplanted larvae were sorted for larvae, which only show tumor cells in the PVS at 2 hours post injection (hpi) and subsequently kept at 34°C.

### Immunoblotting

Protein extraction and immunoblotting was performed according to standard procedures with primary antibodies against FLI1 (1:500, rabbit, ab15289, abcam, UK) and GAPDH (1:5000, rabbit, 5174S, Cell Signaling, UK) and secondary antibody α-rabbit (1:5000, goat anti-rabbit IgG DyLight800 SA5-35571, Invitrogen, USA). Signal detection was performed with the LI-COR Odyssey Infrared Imaging System (LI-COR Bioscience, USA).

### Whole mount immunostaining

Xenotransplanted larvae were fixed in 4% PFA/PBS over night at 4°C. On the following day, larvae were washed once with PBS and then gradually transferred to 100% MetOH to be stored at −20°C until immunostaining. For immunostaining, larvae were gradually transferred back to PBS. After washing with PBSX (PBS with 0.1% Triton X-100) and washing with distilled water, larvae were incubated in acetone (7 min, −20°C). Larvae were blocked for 1 hour in PBDX (PBS with 0.1% Triton X-100, 0.1 g/l BSA, 0.1% DMSO) supplemented with 15 μl goat serum (GS) (normal donor herd, Sigma-Aldrich, USA) per ml PBDX. Primary antibodies against Ki-67 ((8D5) mouse primary mAb #9449, Cell signaling Technology, USA) and Cleaved Caspase 3 ((D175) primary antibody, Cell Signaling Technology, USA) were diluted 1:400 and 1:100 in PBDX+GS, respectively. Larvae were incubated in primary antibody solution over night at 4°C. On the next day, larvae were washed 4x in PBDX. Secondary antibodies Alexa 568 anti-mouse (A-11019, Invitrogen, USA) and Alexa 568 anti-rabbit (A-21069, Invitrogen, USA) were diluted 1:500 in PBDX+GS. Larvae were incubated in secondary antibody solution for 1 hour. From this point on all steps were carried out in the dark. Larvae were washed 2x with PBDX and 1x with PBS, followed by an incubation in 4% PFA for 5 minutes. Larvae were then washed 3x with PBS, transferred to Dako Fluorescence Mounting Medium (Dako, Agilent, USA) and stored at 4°C until imaging.

### Imaging

Larvae were anesthetized in 1x Tricaine. Fluorescent images were acquired using an Axio Zoom.V16 fluorescence stereo zoom microscope with an Axiocam 503 color camera from Zeiss (Zeiss, Germany). Images were acquired using the image software ZEN (Zeiss, Germany). For confocal imaging fixed and immunostained larvae were embedded in 1.2% ultra-low gelling agarose (Cat. No. A2576-25G, Sigma-Aldrich Chemie GmbH, Germany) in a glass bottom imaging dish (D35-14-1.5-NJ, Cellvis, Mountain View, USA) as previously described^43^. Images were acquired using a SP8 X WLL confocal microscope system (Leica, Germany).

### Automated imaging and quantification

For automated imaging, larvae were anesthetized in 1x Tricaine and embedded in a 96-well ZF plate (Hashimoto Electronic Industry Co, Japan) with 0.5 % ultra-low gelling agarose (Cat. No. A2576-25G, Sigma-Aldrich Chemie GmbH, Germany). The Operetta CLS high-content imager (PerkinElmer, USA) was used for image acquisition: 5x air objective, Brightfield (40 ms, 10%), GFP (excitation: 460-490 nm at 100%, emission: 500-550 nm for 400 ms), tagRFP (excitation: 530-560 nm at 100%, emission: 570-650 nm for 400 ms), CellTrace Violet (excitation: 390-420 nm at 100%, emission: 430-500 nm for 600 ms). 23 planes with a distance of 25 μm were imaged per field. Tumor size was quantified with the Harmony Software 4.9 (PerkinElmer, USA). The area of the tumor projected along the z-axis onto the x-y-plane (“footprint area”) was used for further analysis.

### Compound treatment

For drug screening experiments, shSK-E17T Ewing sarcoma cells were transplanted into the PVS of 2 dpf old zebrafish larvae. Tumor cells were allowed to grow for 24 hours to form a compact tumor mass at the injection site. Xenotransplanted larvae were imaged at 1 dpi in the Operetta CLS and then transferred to fresh E3 embryo medium containing respective compounds at NOEC concentrations. All used compounds were purchased from MedChemExpress (Sweden):

**Table 1:**
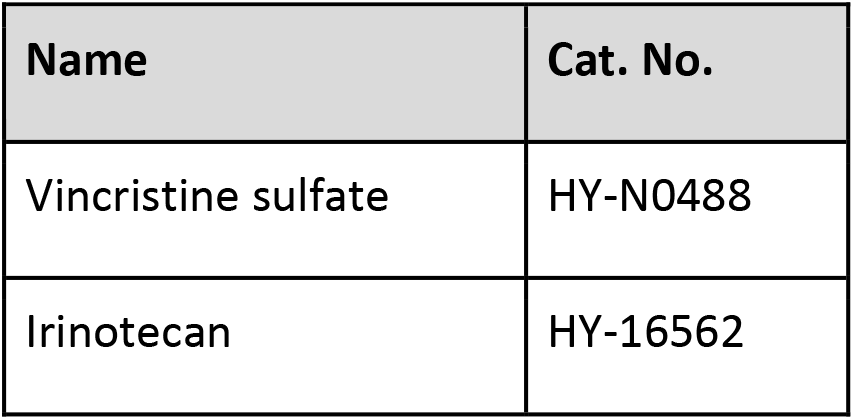
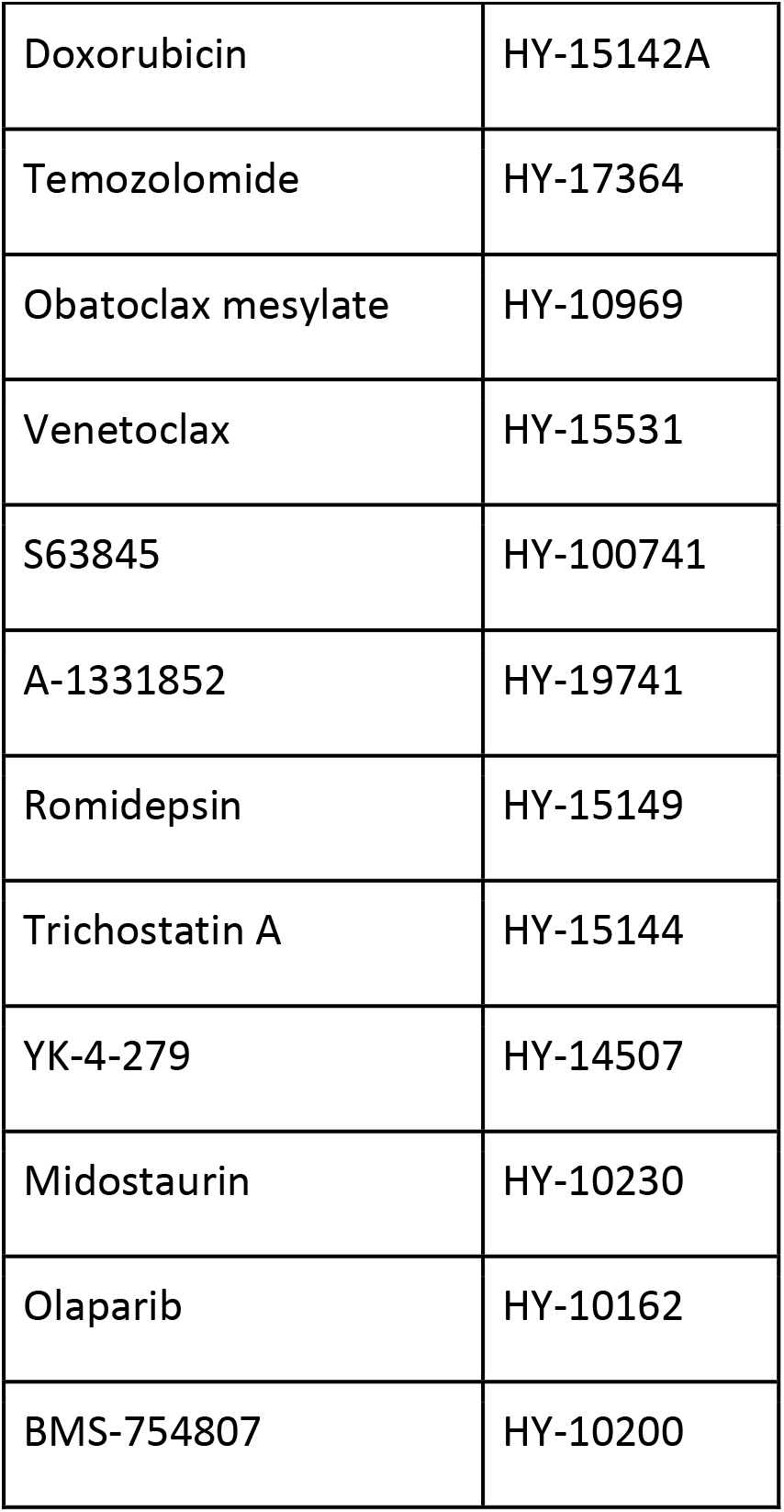
Catalog numbers of compounds.

Compound treatment was performed for 48 hours. At 3 dpi larvae were imaged again post-treatment using the Operetta CLS.

### Gene expression analysis

Pre-processed and normalized gene expression data from cancer cell lines (Cell Line Compendium v2 [December 2019]) and patient samples (Tumor Compendium v11 Public PolyA [April 2020]) were obtained from the Treehouse Childhood Cancer initiative^23^ and processed using R (v 3.6.3) using standard function and the *pheatmap* package. We selected all Ewing-sarcoma-related samples by searching the associated metadata (columns: *“disease”, “Histology”, “Source sample ID”*). Only samples from data sources with at least two Ewing sarcoma samples were considered.

### Statistical analysis

Statistical analysis was performed with Prism 8 (Version 8.3.0, Graphpad, USA). Normal distribution of data was tested with a D’Agostino & Pearson normality test. According to this either parametric tests (student’s t-test or ANOVA) or non-parametric tests (Mann-Whitney test or Kruskal-Wallis test) were performed. The applied analysis method is always indicated in the figure legends of the respective graphs.

## Author contributions

SG designed, performed and analysed experiments and wrote the manuscript, CS, ES, AWW, SP & LB performed and analysed experiments, MT and ET provided the modified A673 cell line and input to the manuscript, DS and OD provided cell lines, patient-derived tumor samples and contributed to writing the manuscript, HK contributed to the design of experiments and to writing the manuscript, FH performed bioinformatic analysis and contributed to writing the manuscript, MD conceptualized the project, designed and analysed experiments and wrote the manuscript.

## Acknowledgement

This work was supported by the Austrian Research Promotion Agency (FFG) project 7940628 (Danio4Can)(M.D.), Alex’s Lemonade Stand Foundation (H.K., F.H. & M.D.) and a DOC fellowship of the Austrian Academy of Sciences to S.G. at St. Anna Children’s Cancer Research Institute. We would like to thank Achim Kirsch for fantastic support with high-content imaging and Karin Mühlbacher for help with cell culture.

## Competing interests

The authors declare to have no competing interests

## Notes

### Competing Interest Statement

The authors have declared no competing interest.

## References

1 Gaspar, N., Hawkins, D. S., Dirksen, U., Lewis, I. J., Ferrari, S., Le Deley, M. C., Kovar, H., Grimer, R., Whelan, J., Claude, L., Delattre, O., Paulussen, M., Picci, P., Sundby Hall, K., van den Berg, H., Ladenstein, R., Michon, J., Hjorth, L., Judson, I., Luksch, R., Bernstein, M. L., Marec-Berard, P., Brennan, B., Craft, A. W., Womer, R. B., Juergens, H. & Oberlin, O. Ewing Sarcoma: Current Management and Future Approaches Through Collaboration. J Clin Oncol 33, 3036–3046, doi:10.1200/JCO.2014.59.5256 (2015).

2 Grunewald, T. G. P., Cidre-Aranaz, F., Surdez, D., Tomazou, E. M., de Alava, E., Kovar, H., Sorensen, P. H., Delattre, O. & Dirksen, U. Ewing sarcoma. Nat Rev Dis Primers 4, 5, doi:10.1038/s41572-018-0003-x (2018).

3 Delattre, O., Zucman, J., Plougastel, B., Desmaze, C., Melot, T., Peter, M., Kovar, H., Joubert, I., de Jong, P., Rouleau, G. & et al. Gene fusion with an ETS DNA-binding domain caused by chromosome translocation in human tumours. Nature 359, 162–165, doi:10.1038/359162a0 (1992).

4 Minas, T. Z., Surdez, D., Javaheri, T., Tanaka, M., Howarth, M., Kang, H. J., Han, J., Han, Z. Y., Sax, B., Kream, B. E., Hong, S. H., Celik, H., Tirode, F., Tuckermann, J., Toretsky, J. A., Kenner, L., Kovar, H., Lee, S., Sweet-Cordero, E. A., Nakamura, T., Moriggl, R., Delattre, O. & Uren, A. Combined experience of six independent laboratories attempting to create an Ewing sarcoma mouse model. Oncotarget 8, 34141–34163, doi:10.18632/oncotarget.9388 (2017).

5 Bierbaumer, L., Katschnig, A. M., Radic-Sarikas, B., Kauer, M. O., Petro, J. A., Hogler, S., Gurnhofer, E., Pedot, G., Schafer, B. W., Schwentner, R., Muhlbacher, K., Kromp, F., Aryee, D. N. T., Kenner, L., Uren, A. & Kovar, H. YAP/TAZ inhibition reduces metastatic potential of Ewing sarcoma cells. Oncogenesis 10, 2, doi:10.1038/s41389-020-00294-8 (2021).

6 Iniguez, A. B., Stolte, B., Wang, E. J., Conway, A. S., Alexe, G., Dharia, N. V., Kwiatkowski, N., Zhang, T., Abraham, B. J., Mora, J., Kalev, P., Leggett, A., Chowdhury, D., Benes, C. H., Young, R. A., Gray, N. S. & Stegmaier, K. EWS/FLI Confers Tumor Cell Synthetic Lethality to CDK12 Inhibition in Ewing Sarcoma. Cancer Cell 33, 202–216 e206, doi:10.1016/j.ccell.2017.12.009 (2018).

7 Surdez, D., Landuzzi, L., Scotlandi, K. & Manara, M. C. Ewing Sarcoma PDX Models. Methods Mol Biol 2226, 223–242, doi:10.1007/978-1-0716-1020-6_18 (2021).

8 Aynaud, M. M., Mirabeau, O., Gruel, N., Grossetete, S., Boeva, V., Durand, S., Surdez, D., Saulnier, O., Zaidi, S., Gribkova, S., Fouche, A., Kairov, U., Raynal, V., Tirode, F., Grunewald, T. G. P., Bohec, M., Baulande, S., Janoueix-Lerosey, I., Vert, J. P., Barillot, E., Delattre, O. & Zinovyev, A. Transcriptional Programs Define Intratumoral Heterogeneity of Ewing Sarcoma at Single-Cell Resolution. Cell Rep 30, 1767–1779 e1766, doi:10.1016/j.celrep.2020.01.049 (2020).

9 Fior, R., Povoa, V., Mendes, R. V., Carvalho, T., Gomes, A., Figueiredo, N. & Ferreira, M. G. Single-cell functional and chemosensitive profiling of combinatorial colorectal therapy in zebrafish xenografts. Proc Natl Acad Sci U S A 114, E8234–E8243, doi:10.1073/pnas.1618389114 (2017).

10 Lee, L. M., Seftor, E. A., Bonde, G., Cornell, R. A. & Hendrix, M. J. The fate of human malignant melanoma cells transplanted into zebrafish embryos: assessment of migration and cell division in the absence of tumor formation. Dev Dyn 233, 1560–1570, doi:10.1002/dvdy.20471 (2005).

11 Nicoli, S. & Presta, M. The zebrafish/tumor xenograft angiogenesis assay. Nat Protoc 2, 2918–2923, doi:10.1038/nprot.2007.412 (2007).

12 Veinotte, C. J., Dellaire, G. & Berman, J. N. Hooking the big one: the potential of zebrafish xenotransplantation to reform cancer drug screening in the genomic era. Dis Model Mech 7, 745–754, doi:10.1242/dmm.015784 (2014).

13 Yan, C., Brunson, D. C., Tang, Q., Do, D., Iftimia, N. A., Moore, J. C., Hayes, M. N., Welker, A. M., Garcia, E. G., Dubash, T. D., Hong, X., Drapkin, B. J., Myers, D. T., Phat, S., Volorio, A., Marvin, D. L., Ligorio, M., Dershowitz, L., McCarthy, K. M., Karabacak, M. N., Fletcher, J. A., Sgroi, D. C., Iafrate, J. A., Maheswaran, S., Dyson, N. J., Haber, D. A., Rawls, J. F. & Langenau, D. M. Visualizing Engrafted Human Cancer and Therapy Responses in Immunodeficient Zebrafish. Cell 177, 1903–1914 e1914, doi:10.1016/j.cell.2019.04.004 (2019).

14 Barriuso, J., Nagaraju, R. & Hurlstone, A. Zebrafish: a new companion for translational research in oncology. Clin Cancer Res 21, 969–975, doi:10.1158/1078-0432.CCR-14-2921 (2015).

15 Cully, M. Zebrafish earn their drug discovery stripes. Nat Rev Drug Discov 18, 811–813, doi:10.1038/d41573-019-00165-x (2019).

16 Rennekamp, A. J. & Peterson, R. T. 15 years of zebrafish chemical screening. Curr Opin Chem Biol 24, 58–70, doi:10.1016/j.cbpa.2014.10.025 (2015).

17 Ban, J., Aryee, D. N., Fourtouna, A., van der Ent, W., Kauer, M., Niedan, S., Machado, I., Rodriguez-Galindo, C., Tirado, O. M., Schwentner, R., Picci, P., Flanagan, A. M., Berg, V., Strauss, S. J., Scotlandi, K., Lawlor, E. R., Snaar-Jagalska, E., Llombart-Bosch, A. & Kovar, H. Suppression of deacetylase SIRT1 mediates tumor-suppressive NOTCH response and offers a novel treatment option in metastatic Ewing sarcoma. Cancer Res 74, 6578–6588, doi:10.1158/0008-5472.CAN-14-1736 (2014).

18 van der Ent, W., Jochemsen, A. G., Teunisse, A. F., Krens, S. F., Szuhai, K., Spaink, H. P., Hogendoorn, P. C. & Snaar-Jagalska, B. E. Ewing sarcoma inhibition by disruption of EWSR1-FLI1 transcriptional activity and reactivation of p53. J Pathol 233, 415–424, doi:10.1002/path.4378 (2014).

19 Franzetti, G. A., Laud-Duval, K., van der Ent, W., Brisac, A., Irondelle, M., Aubert, S., Dirksen, U., Bouvier, C., de Pinieux, G., Snaar-Jagalska, E., Chavrier, P. & Delattre, O. Cell-to-cell heterogeneity of EWSR1-FLI1 activity determines proliferation/migration choices in Ewing sarcoma cells. Oncogene 36, 3505–3514, doi:10.1038/onc.2016.498 (2017).

20 Barber-Rotenberg, J. S., Selvanathan, S. P., Kong, Y., Erkizan, H. V., Snyder, T. M., Hong, S. P., Kobs, C. L., South, N. L., Summer, S., Monroe, P. J., Chruszcz, M., Dobrev, V., Tosso, P. N., Scher, L. J., Minor, W., Brown, M. L., Metallo, S. J., Uren, A. & Toretsky, J. A. Single enantiomer of YK-4-279 demonstrates specificity in targeting the oncogene EWS-FLI1. Oncotarget 3, 172–182, doi:10.18632/oncotarget.454 (2012).

21 Erkizan, H. V., Kong, Y., Merchant, M., Schlottmann, S., Barber-Rotenberg, J. S., Yuan, L., Abaan, O. D., Chou, T. H., Dakshanamurthy, S., Brown, M. L., Uren, A. & Toretsky, J. A. A small molecule blocking oncogenic protein EWS-FLI1 interaction with RNA helicase A inhibits growth of Ewing’s sarcoma. Nat Med 15, 750–756, doi:10.1038/nm.1983 (2009).

22 Katschnig, A. M., Kauer, M. O., Schwentner, R., Tomazou, E. M., Mutz, C. N., Linder, M., Sibilia, M., Alonso, J., Aryee, D. N. T. & Kovar, H. EWS-FLI1 perturbs MRTFB/YAP-1/TEAD target gene regulation inhibiting cytoskeletal autoregulatory feedback in Ewing sarcoma. Oncogene 36, 5995–6005, doi:10.1038/onc.2017.202 (2017).

23 Bjork, I., Peralez, J., Haussler, D., Spunt, S. L. & Vaske, O. M. Data sharing for clinical utility. Cold Spring Harb Mol Case Stud 5, doi:10.1101/mcs.a004689 (2019).

24 Xiao, J., Glasgow, E. & Agarwal, S. Zebrafish Xenografts for Drug Discovery and Personalized Medicine. Trends Cancer 6, 569–579, doi:10.1016/j.trecan.2020.03.012 (2020).

25 Zhang, B., Shimada, Y., Kuroyanagi, J., Umemoto, N., Nishimura, Y. & Tanaka, T. Quantitative phenotyping-based in vivo chemical screening in a zebrafish model of leukemia stem cell xenotransplantation. PLoS One 9, e85439, doi:10.1371/journal.pone.0085439 (2014).

26 Palmerini, E., Jones, R. L., Setola, E., Picci, P., Marchesi, E., Luksch, R., Grignani, G., Cesari, M., Longhi, A., Abate, M. E., Paioli, A., Szucs, Z., D’Ambrosio, L., Scotlandi, K., Fagioli, F., Asaftei, S. & Ferrari, S. Irinotecan and temozolomide in recurrent Ewing sarcoma: an analysis in 51 adult and pediatric patients. Acta Oncol 57, 958–964, doi:10.1080/0284186X.2018.1449250 (2018).

27 Kehr, S., Haydn, T., Bierbrauer, A., Irmer, B., Vogler, M. & Fulda, S. Targeting BCL-2 proteins in pediatric cancer: Dual inhibition of BCL-XL and MCL-1 leads to rapid induction of intrinsic apoptosis. Cancer Lett 482, 19–32, doi:10.1016/j.canlet.2020.02.041 (2020).

28 Roberts, A. W., Davids, M. S., Pagel, J. M., Kahl, B. S., Puvvada, S. D., Gerecitano, J. F., Kipps, T. J., Anderson, M. A., Brown, J. R., Gressick, L., Wong, S., Dunbar, M., Zhu, M., Desai, M. B., Cerri, E., Heitner Enschede, S., Humerickhouse, R. A., Wierda, W. G. & Seymour, J. F. Targeting BCL2 with Venetoclax in Relapsed Chronic Lymphocytic Leukemia. N Engl J Med 374, 311–322, doi:10.1056/NEJMoa1513257 (2016).

29 Montero, J. & Letai, A. Why do BCL-2 inhibitors work and where should we use them in the clinic? Cell Death Differ 25, 56–64, doi:10.1038/cdd.2017.183 (2018).

30 Dharia, N. V., Kugener, G., Guenther, L. M., Malone, C. F., Durbin, A. D., Hong, A. L., Howard, T. P., Bandopadhayay, P., Wechsler, C. S., Fung, I., Warren, A. C., Dempster, J. M., Krill-Burger, J. M., Paolella, B. R., Moh, P., Jha, N., Tang, A., Montgomery, P., Boehm, J. S., Hahn, W. C., Roberts, C. W. M., McFarland, J. M., Tsherniak, A., Golub, T. R., Vazquez, F. & Stegmaier, K. A first-generation pediatric cancer dependency map. Nat Genet, doi:10.1038/s41588-021-00819-w (2021).

31 Tsherniak, A., Vazquez, F., Montgomery, P. G., Weir, B. A., Kryukov, G., Cowley, G. S., Gill, S., Harrington, W. F., Pantel, S., Krill-Burger, J. M., Meyers, R. M., Ali, L., Goodale, A., Lee, Y., Jiang, G., Hsiao, J., Gerath, W. F. J., Howell, S., Merkel, E., Ghandi, M., Garraway, L. A., Root, D. E., Golub, T. R., Boehm, J. S. & Hahn, W. C. Defining a Cancer Dependency Map. Cell 170, 564–576 e516, doi:10.1016/j.cell.2017.06.010 (2017).

32 Wang, L., Doherty, G. A., Judd, A. S., Tao, Z. F., Hansen, T. M., Frey, R. R., Song, X., Bruncko, M., Kunzer, A. R., Wang, X., Wendt, M. D., Flygare, J. A., Catron, N. D., Judge, R. A., Park, C. H., Shekhar, S., Phillips, D. C., Nimmer, P., Smith, M. L., Tahir, S. K., Xiao, Y., Xue, J., Zhang, H., Le, P. N., Mitten, M. J., Boghaert, E. R., Gao, W., Kovar, P., Choo, E. F., Diaz, D., Fairbrother, W. J., Elmore, S. W., Sampath, D., Leverson, J. D. & Souers, A. J. Discovery of A-1331852, a First-in-Class, Potent, and Orally-Bioavailable BCL-XL Inhibitor. ACS Med Chem Lett 11, 1829–1836, doi:10.1021/acsmedchemlett.9b00568 (2020).

33 Bolomsky, A., Vogler, M., Kose, M. C., Heckman, C. A., Ehx, G., Ludwig, H. & Caers, J. MCL-1 inhibitors, fast-lane development of a new class of anti-cancer agents. J Hematol Oncol 13, 173, doi:10.1186/s13045-020-01007-9 (2020).

34 Mukherjee, N., Skees, J., Todd, K. J., West, D. A., Lambert, K. A., Robinson, W. A., Amato, C. M., Couts, K. L., Van Gulick, R., MacBeth, M., Nassar, K., Tan, A. C., Zhai, Z., Fujita, M., Bagby, S. M., Dart, C. R., Lambert, J. R., Norris, D. A. & Shellman, Y. G. MCL1 inhibitors S63845/MIK665 plus Navitoclax synergistically kill difficult-to-treat melanoma cells. Cell Death Dis 11, 443, doi:10.1038/s41419-020-2646-2 (2020).

35 Sarosiek, K. A., Fraser, C., Muthalagu, N., Bhola, P. D., Chang, W., McBrayer, S. K., Cantlon, A., Fisch, S., Golomb-Mello, G., Ryan, J. A., Deng, J., Jian, B., Corbett, C., Goldenberg, M., Madsen, J. R., Liao, R., Walsh, D., Sedivy, J., Murphy, D. J., Carrasco, D. R., Robinson, S., Moslehi, J. & Letai, A. Developmental Regulation of Mitochondrial Apoptosis by c-Myc Governs Age- and Tissue-Specific Sensitivity to Cancer Therapeutics. Cancer Cell 31, 142–156, doi:10.1016/j.ccell.2016.11.011 (2017).

36 Mason, K. D., Carpinelli, M. R., Fletcher, J. I., Collinge, J. E., Hilton, A. A., Ellis, S., Kelly, P. N., Ekert, P. G., Metcalf, D., Roberts, A. W., Huang, D. C. & Kile, B. T. Programmed anuclear cell death delimits platelet life span. Cell 128, 1173–1186, doi:10.1016/j.cell.2007.01.037 (2007).

37 Li, Z., He, S. & Look, A. T. The MCL1-specific inhibitor S63845 acts synergistically with venetoclax/ABT-199 to induce apoptosis in T-cell acute lymphoblastic leukemia cells. Leukemia 33, 262–266, doi:10.1038/s41375-018-0201-2 (2019).

38 Carrington, E. M., Zhan, Y., Brady, J. L., Zhang, J. G., Sutherland, R. M., Anstee, N. S., Schenk, R. L., Vikstrom, I. B., Delconte, R. B., Segal, D., Huntington, N. D., Bouillet, P., Tarlinton, D. M., Huang, D. C., Strasser, A., Cory, S., Herold, M. J. & Lew, A. M. Anti-apoptotic proteins BCL-2, MCL-1 and A1 summate collectively to maintain survival of immune cell populations both in vitro and in vivo. Cell Death Differ 24, 878–888, doi:10.1038/cdd.2017.30 (2017).

39 Kimmel, C. B., Ballard, W. W., Kimmel, S. R., Ullmann, B. & Schilling, T. F. Stages of embryonic development of the zebrafish. Dev Dyn 203, 253–310, doi:10.1002/aja.1002030302 (1995).

40 Westerfield, M. The zebrafish book. A Guide for the Laboratory Use of Zebrafish (Danio rerio). The University of Oregon press 4th edition(2000).

41 Stewart, E., Federico, S. M., Chen, X., Shelat, A. A., Bradley, C., Gordon, B., Karlstrom, A., Twarog, N. R., Clay, M. R., Bahrami, A., Freeman, B. B., 3rd, Xu, B., Zhou, X., Wu, J., Honnell, V., Ocarz, M., Blankenship, K., Dapper, J., Mardis, E. R., Wilson, R. K., Downing, J., Zhang, J., Easton, J., Pappo, A. & Dyer, M. A. Orthotopic patient-derived xenografts of paediatric solid tumours. Nature 549, 96–100, doi:10.1038/nature23647 (2017).

42 Pascoal, S., Grissenberger, S., Scheuringer, E., Fior, R., Ferreira, M. G. & Distel, M. Using Zebrafish Larvae as a Xenotransplantation Model to Study Ewing Sarcoma. Methods Mol Biol 2226, 243–255, doi:10.1007/978-1-0716-1020-6_19 (2021).

43 Distel, M. & Koster, R. W. In vivo time-lapse imaging of zebrafish embryonic development. CSH Protoc 2007, pdb prot4816, doi:10.1101/pdb.prot4816 (2007).

44 Tsafou, K., Katschnig, A. M., Radic-Sarikas, B., Mutz, C. N., Iljin, K., Schwentner, R., Kauer, M. O., Muhlbacher, K., Aryee, D. N. T., Westergaard, D., Haapa-Paananen, S., Fey, V., Superti-Furga, G., Toretsky, J., Brunak, S. & Kovar, H. Identifying the druggable interactome of EWS-FLI1 reveals MCL-1 dependent differential sensitivities of Ewing sarcoma cells to apoptosis inducers. Oncotarget 9, 31018–31031, doi:10.18632/oncotarget.25760 (2018).

45 Heisey, D. A. R., Lochmann, T. L., Floros, K. V., Coon, C. M., Powell, K. M., Jacob, S., Calbert, M. L., Ghotra, M. S., Stein, G. T., Maves, Y. K., Smith, S. C., Benes, C. H., Leverson, J. D., Souers, A. J., Boikos, S. A. & Faber, A. C. The Ewing Family of Tumors Relies on BCL-2 and BCL-XL to Escape PARP Inhibitor Toxicity. Clin Cancer Res 25, 1664–1675, doi:10.1158/1078-0432.CCR-18-0277 (2019).

46 Sakimura, R., Tanaka, K., Nakatani, F., Matsunobu, T., Li, X., Hanada, M., Okada, T., Nakamura, T., Matsumoto, Y. & Iwamoto, Y. Antitumor effects of histone deacetylase inhibitor on Ewing’s family tumors. Int J Cancer 116, 784–792, doi:10.1002/ijc.21069 (2005).

47 Souza, B. K., da Costa Lopez, P. L., Menegotto, P. R., Vieira, I. A., Kersting, N., Abujamra, A. L., Brunetto, A. T., Brunetto, A. L., Gregianin, L., de Farias, C. B., Thiele, C. J. & Roesler, R. Targeting Histone Deacetylase Activity to Arrest Cell Growth and Promote Neural Differentiation in Ewing Sarcoma. Mol Neurobiol 55, 7242–7258, doi:10.1007/s12035-018-0874-6 (2018).

48 Boro, A., Pretre, K., Rechfeld, F., Thalhammer, V., Oesch, S., Wachtel, M., Schafer, B. W. & Niggli, F. K. Small-molecule screen identifies modulators of EWS/FLI1 target gene expression and cell survival in Ewing’s sarcoma. Int J Cancer 131, 2153–2164, doi:10.1002/ijc.27472 (2012).

49 Brenner, J. C., Feng, F. Y., Han, S., Patel, S., Goyal, S. V., Bou-Maroun, L. M., Liu, M., Lonigro, R., Prensner, J. R., Tomlins, S. A. & Chinnaiyan, A. M. PARP-1 inhibition as a targeted strategy to treat Ewing’s sarcoma. Cancer Res 72, 1608–1613, doi:10.1158/0008-5472.CAN-11-3648 (2012).

50 Garnett, M. J., Edelman, E. J., Heidorn, S. J., Greenman, C. D., Dastur, A., Lau, K. W., Greninger, P., Thompson, I. R., Luo, X., Soares, J., Liu, Q., Iorio, F., Surdez, D., Chen, L., Milano, R. J., Bignell, G. R., Tam, A. T., Davies, H., Stevenson, J. A., Barthorpe, S., Lutz, S. R., Kogera, F., Lawrence, K., McLaren-Douglas, A., Mitropoulos, X., Mironenko, T., Thi, H., Richardson, L., Zhou, W., Jewitt, F., Zhang, T., O’Brien, P., Boisvert, J. L., Price, S., Hur, W., Yang, W., Deng, X., Butler, A., Choi, H. G., Chang, J. W., Baselga, J., Stamenkovic, I., Engelman, J. A., Sharma, S. V., Delattre, O., Saez-Rodriguez, J., Gray, N. S., Settleman, J., Futreal, P. A., Haber, D. A., Stratton, M. R., Ramaswamy, S., McDermott, U. & Benes, C. H. Systematic identification of genomic markers of drug sensitivity in cancer cells. Nature 483, 570–575, doi:10.1038/nature11005 (2012).

51 Engert, F., Schneider, C., Weibeta, L. M., Probst, M. & Fulda, S. PARP Inhibitors Sensitize Ewing Sarcoma Cells to Temozolomide-Induced Apoptosis via the Mitochondrial Pathway. Mol Cancer Ther 14, 2818–2830, doi:10.1158/1535-7163.MCT-15-0587 (2015).

52 Bailey, K., Cost, C., Davis, I., Glade-Bender, J., Grohar, P., Houghton, P., Isakoff, M., Stewart, E., Laack, N., Yustein, J., Reed, D., Janeway, K., Gorlick, R., Lessnick, S., DuBois, S. & Hingorani, P. Emerging novel agents for patients with advanced Ewing sarcoma: a report from the Children’s Oncology Group (COG) New Agents for Ewing Sarcoma Task Force. F1000Res 8, doi:10.12688/f1000research.18139.1 (2019).

53 Celik, H., Sciandra, M., Flashner, B., Gelmez, E., Kayraklioglu, N., Allegakoen, D. V., Petro, J. R., Conn, E. J., Hour, S., Han, J., Oktay, L., Tiwari, P. B., Hayran, M., Harris, B. T., Manara, M. C., Toretsky, J. A., Scotlandi, K. & Uren, A. Clofarabine inhibits Ewing sarcoma growth through a novel molecular mechanism involving direct binding to CD99. Oncogene 37, 2181–2196, doi:10.1038/s41388-017-0080-4 (2018).

