## Supplemental Figures and Tables for "Automated compound testing in zebrafish xenografts identifies combined MCL-1 and BCL-X_L_ inhibition to be effective against Ewing sarcoma"

### Supplementary Figures & Tables

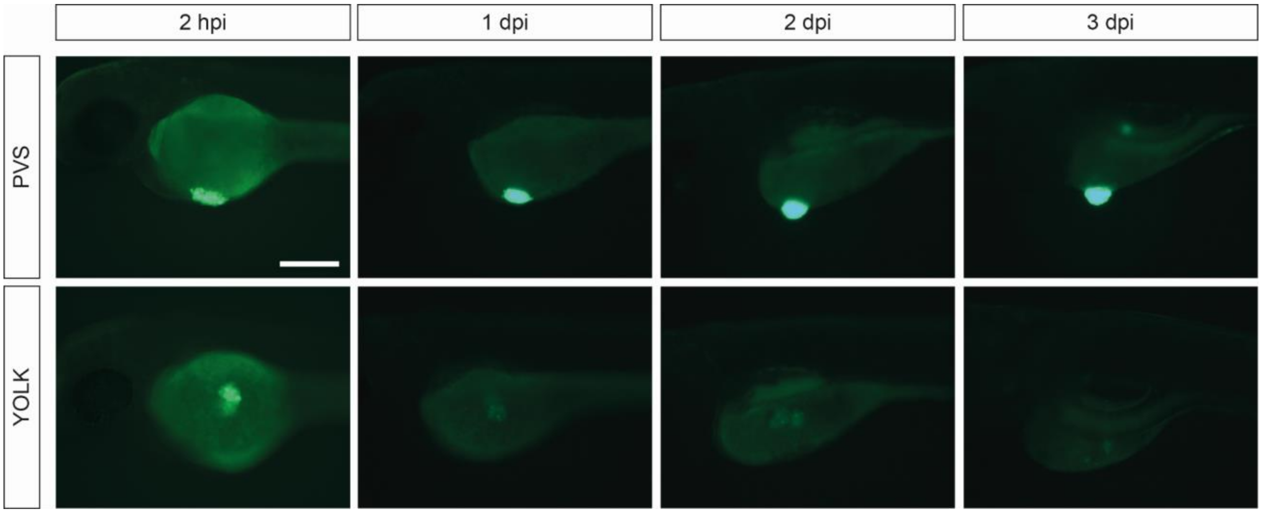


**Figure S1**: **Comparison of injection sites**
2 dpf old zebrafish embryos were transplanted with shSK-E17T Ewing sarcoma cells either in the perivitelline space (PVS) (upper panel) or into the yolk (lower panel) and imaged at different time points (2 hpi, 1 dpi, 2 dpi, 3 dpi). Scale bar is 250 µm.

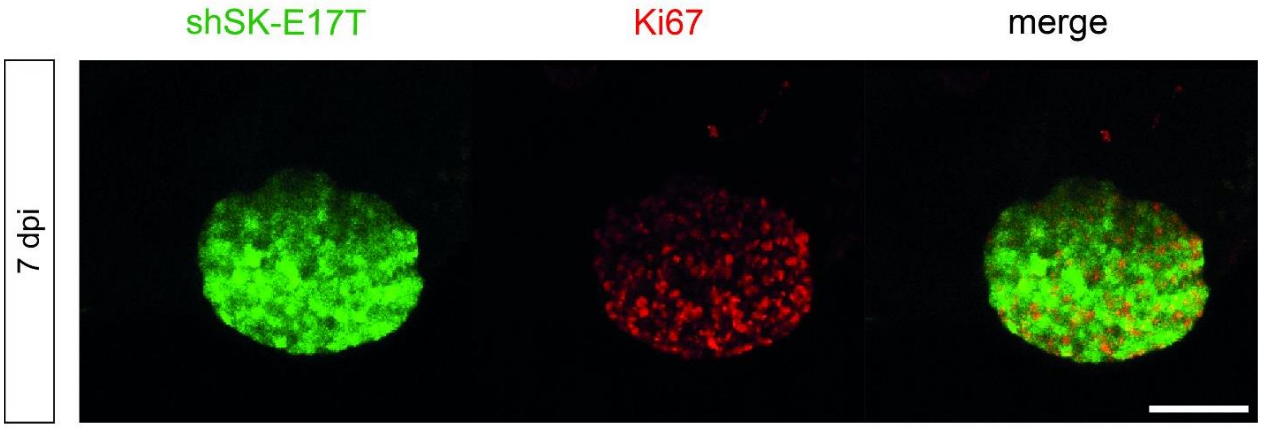


**Figure S2**: **Ki67 immunostaining at 7 dpi**
2 dpf old zebrafish embryo was transplanted with shSK-E17T Ewing sarcoma cells, fixed at 7 dpi and subsequently immunostained with a human-specific Ki67 antibody. Scale bar is 100 µm.

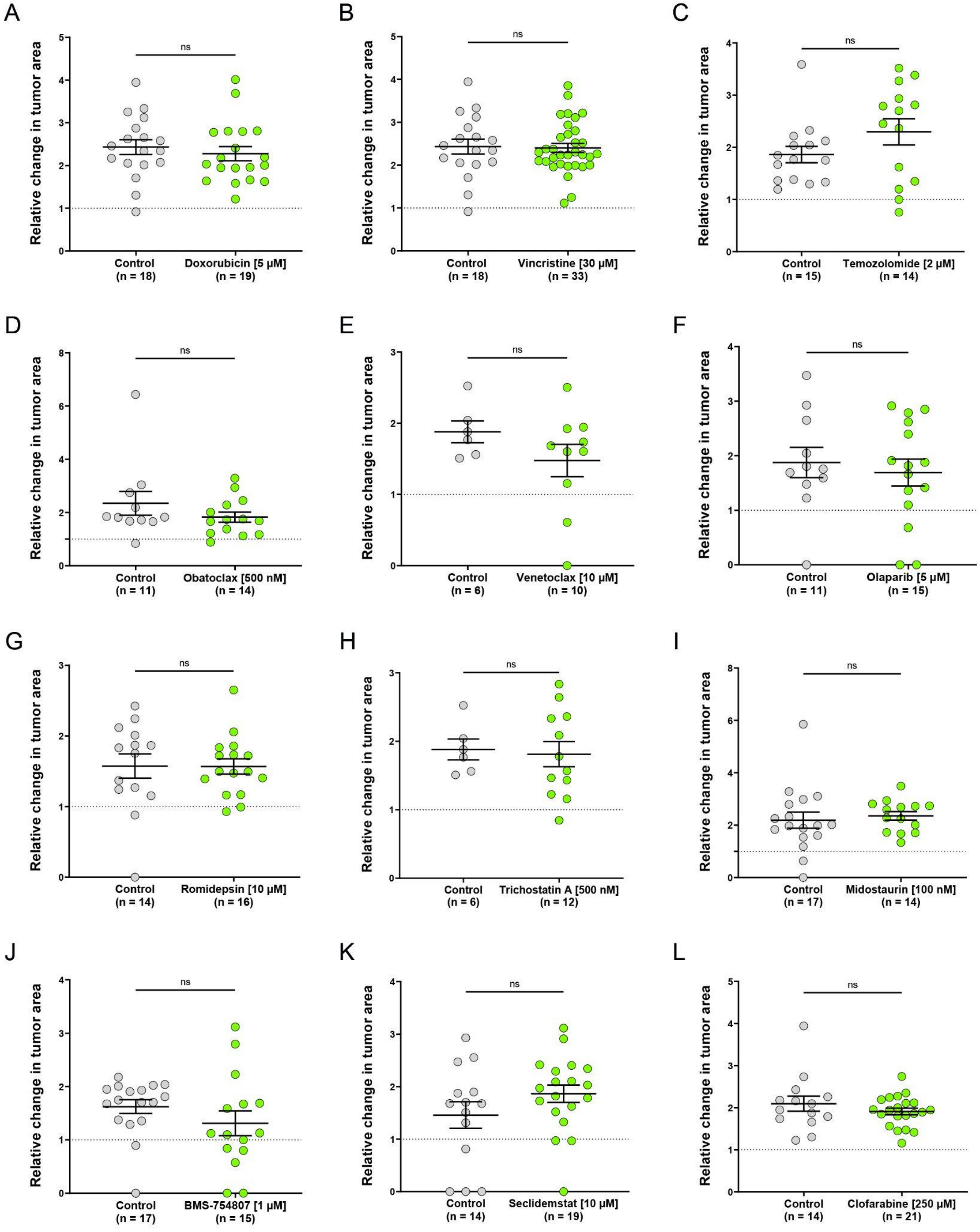


**Figure S3**: **single compound treatment of shSK-E17T zebrafish xenografts**
Xenotransplanted larvae were treated with Doxorubicin (5 µM, A), Vincristine (30 µM, B), Temozolomide (2 µM, C), Obatoclax (500 nM, D), Venetoclax (10 µM, E), Olaparib (5 µM, F), Romidepsin (10 µM, G), Tricostatin A (500 nM, H), Midostaurin (100 nM, I), BMS-754807 (1 µM, J), Seclidemstat (10 µM, K) and clofarabine (250 µM, L) for 48 hours. Dot plots show relative change of tumor area (3 dpi/1 dpi). Statistical analyses were performed with a Mann-Whitney test (C, D, E, H, I, J, L) or an unpaired student's t-test (A, B, F, G, K). Error bars represent SEM of n individual larvae.

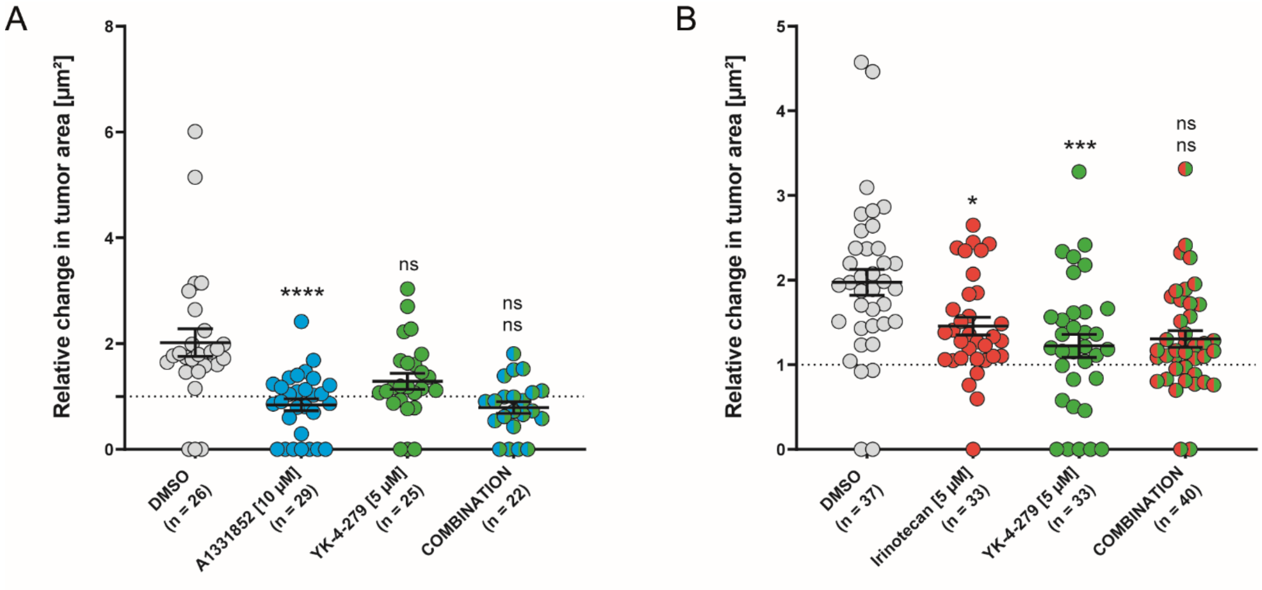


**Figure S4: Non-effective combination treatments in shSK-E17T-transplanted embryos**
Xenotransplanted larvae were treated with A-1331852 [10 µM], YK-4-279 [5 µM] and combination (A) and irinotecan [5 µM], YK-4-279 [5 µM] and combination (B) for 48 hours (1 dpi - 3 dpi). Dot plot shows relative change of tumor area (3 dpi/1 dpi). Significance is shown as followed: Single treatments were compared to DMSO control. Combination treatments were compared to respective single treatments (top value indicates comparison to first compound, bottom value indicates comparison to second compound). Statistical analyses were performed with a Kruskal-Wallis test, ****: $p \leq 0.0001$, ***: $p \leq 0.001$, *: $p \leq 0.05$. Error bars represent SEM of combined larvae (n) of two individual experiments.

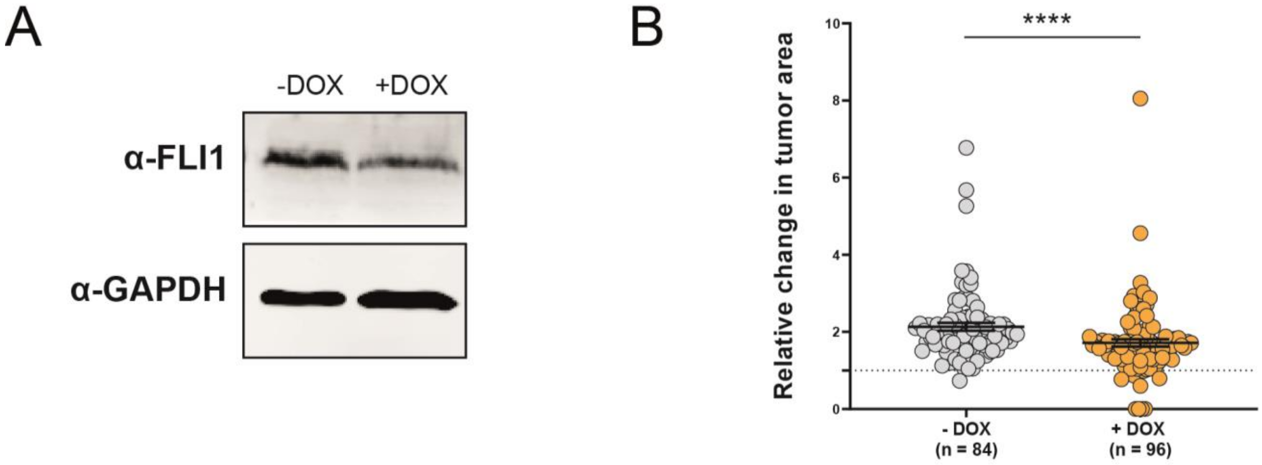


**Figure S5**: **EWS-FLI1 knock down**

A) shSK-E17T Ewing sarcoma cells were treated with 1 µg/ml Doxycycline (DOX) for 7 days to induce shRNA-dependent knock down of EWS-FLI1. To confirm this downregulation a western blot analysis was performed with antibodies against FLI1 and GAPDH. B) Zebrafish embryos were transplanted with either untreated or DOX-pre-treated (>7 days) shSK-E17T cells. Embryos were imaged at 1 dpi and 3 dpi and the relative tumor size was calculated (3 dpi/1 dpi). Statistical analysis was performed with a Mann-Whitney test, ****: p≤0.0001. Error bars represent SEM of combined larvae (n) of 5 independent experiments

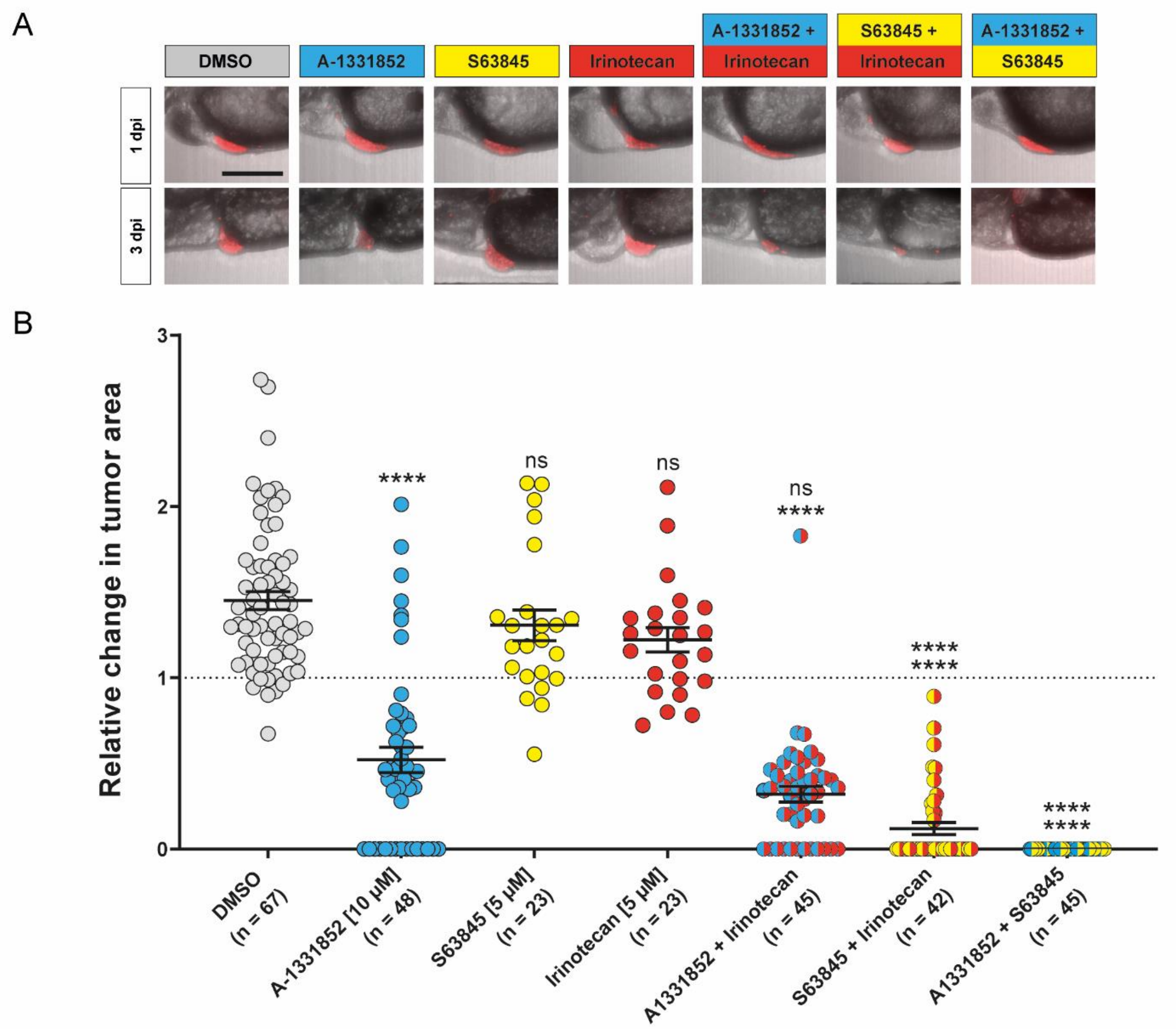


C

| | mean absolute tumor size [µm²] | | % mean increase/decrease |
|---|---|---|---|
| | 1 dpi | 3 dpi | |
| DMSO | 7713 | 10538 | 36,6 |
| A-1331852 [10 µM] | 7116 | 3586 | -49,6 |
| S63845 [5 µM] | 7596 | 9182 | 20,9 |
| Irinotecan [5 µM] | 7271 | 8623 | 18,6 |
| A-1331852 + Irinotecan | 7957 | 2565 | -67,8 |
| S63845 + Irinotecan | 7502 | 648 | -91,4 |
| A-1331852 + S63845 | 7406 | 0 | -100,0 |

**Figure S6: Combination treatments in A673-1c-transplanted embryos**

Xenotransplanted larvae were treated with A-1331852 (10 µM), S63845 (5 µM), Irinotecan (5 µM), and respective combinations for 48 hours (1 dpi - 3 dpi). A) Representative images for average change in tumor size for every condition. B) Relative change of tumor area (3 dpi/1 dpi). C) Mean average size and change in percentage.

Scale bar is 250 µm. Significance is shown as followed: Single treatments were compared to DMSO control. Combination treatments were compared to respective single treatments (top value indicates comparison to first compound, bottom value indicates comparison to second compound). Statistical analyses were performed with a Kruskal-Wallis test, ****: p≤0.0001, **: p≤0.01, *: p≤0.05. Error bars represent SEM of combined larvae (n) of one experiment (single treatments) or two independent experiments (combinations).

**Table S1: Compounds used in our *in vivo* study**

The determined “no observed effect concentration” (NOEC) in zebrafish is stated on the right.

| Compound | Biological function | Reference | NOEC |
|---|---|---|---|
| Vincristine | Microtubule inhibitor | Approved or in clinical trials for Ewing sarcoma treatment | >30 µM |
| Irinotecan | Topoisomerase I inhibitor | | >100 µM |
| Doxorubicin | Topoisomerase II inhibitor | | 5 µM |
| Temozolomide | Alkylating agent | Alternative to Ifosfamide | 2 µM |
| Obatoclax | Pan-Bcl-2 family inhibitor | [44] | 500 nM |
| Venetoclax | BCL-2 inhibitor | [45] together with PARPi | 10 µM |
| S63845 | MCL-1 inhibitor | [44] | 10 µM |
| A-1331852 | BCL-$X_L$ inhibitor | [27] | 10 µM |
| Romidepsin | HDAC inhibitor | [46] | 10 µM |
| Trichostatin A | HDAC inhibitor | [47] | 500 nM |
| YK-4-279 | Inhibitor of EWS-FLI1-RNA helicase A interaction | [21] | 10 µM |
| Midostaurin | Kinase inhibitor | [48] | 100 nM |
| Olaparib | PARP inhibitor | [49-51] | 5 µM |
| BMS-754807 | IGF1R inhibitor | [52] | 1 µM |
| Clofarabine | CD99 inhibitor | [53] | >50 µM |
| Seclidemstat | LSD1 inhibitor | [52] | 10 µM |

**Table S2: NOECs for combination treatments**

| Combination | NOEC |
|---|---|
| S63845 + YK-4-279 | 5 µM + 5 µM |
| S63845 + A-1331852 | 5 µM + 10 µM |
| S63845 + Irinotecan | 5 µM + 5 µM |
| A-1331852 + YK-4-279 | 10 µM + 5 µM |
| A-1331852 + Irinotecan | 10 µM + 5 µM |
| Irinotecan + YK-4-279 | 5 µM + 5 µM |
